# PV - Oligodendrocyte Interactions in the Infralimbic Cortex Promote Extracellular Plasticity after Safety Learning

**DOI:** 10.1101/2025.07.25.666881

**Authors:** L.E. Denholtz, J. Liu, I. Nahmoud, P. Casaccia, E. Likhtik

**Affiliations:** Biology Department, Hunter College, CUNY; Biology Program, The Graduate Center, CUNY; Advanced Science Research Center, CUNY; Department of Psychiatry and Behavioral Neuroscience, Wayne State University School of Medicine

**Keywords:** oligodendrocytes, satellite cells, maturation stage, perineuronal nets, parvalbumin cells, infralimbic, cortex, safety learning, plasticity

## Abstract

Safety learning is mediated by the infralimbic region (IL) of the medial prefrontal cortex, but its cellular mechanisms are poorly understood. Here we show that safety learning improves cognitive flexibility in the long-term, which is associated with recruitment of oligodendrocyte progenitor cells to a satellite position at parvalbumin (PV) interneurons in the IL and their maturation into oligodendrocytes (OLs), as well as a decrease in perineuronal nets (PNNs) surrounding PVs paired with satellite OLs. Using scRNA transcriptomic data mining, we demonstrate that immature OLs primarily express PNN assembly genes, whereas mature OLs express PNN degradation enzymes. We then demonstrate that inhibiting IL PVs during safety learning prevents safety-induced cognitive flexibility, satellite OL maturation, and PNN degradation around IL PV. Thus, we propose that safety learning drives a novel form of neuro-glial plasticity that helps degrade PNNs around PV interneurons via OL recruitment and maturation, thereby shaping IL long-term activity.

---

Identifying stimuli that signal threat or safety is essential to drive avoidance or exploration while navigating changing circumstances. Continuous discrimination of stimulus valence relies on cognitive flexibility, and an imbalance towards overly generalized fear can lead to maladaptive behaviors, characteristic of stress- and anxiety-disorders such as post-traumatic stress disorder (PTSD)^1–4^. Indeed, impairments in cognitive flexibility and poor fear suppression are key behavioral features of PTSD^1,5–7^.

Parvalbumin (PV) interneurons (IN) within the medial prefrontal cortex (mPFC) are essential for local-circuit activity and sculpt oscillatory patterns that set circuit-level communication with other regions during threat-safety discrimination^8–16^. The infralimbic (IL) region of the mPFC is key for processing fear suppression^17–25^, but the role of IL PV activity in emotion regulation is not well understood ^11,26–28^. However, previous work reported that gamma oscillations, which are attributed to PV INs increase in the IL during fear extinction^29,30^, suggesting that PV cells in the IL may contribute to fear suppression behavior. Thus, in this study we asked whether PV IN activity is differentially modulated in the IL during fear and safety learning.

One non-neuronal mechanism for regulating PV activity and plasticity is the perineuronal net (PNN)^31^, a specialized form of condensed extracellular matrix with a honeycomb-like structure that is found primarily around the soma of GABAergic neurons ^32–35^. PNNs form around PV INs during development, decrease with the closure of the critical period (8-10 years old in humans), and mark a phase of brain maturation wherein synapse remodeling is more limited ^36–40^. In adulthood, PNNs are reported to stabilize long-term memory engrams and undergo activity-dependent remodeling during fear learning ^38,41–45^. However, the mechanisms regulating the activity-dependent remodeling of PNN around PV INs remain poorly understood and were investigated in this study.

Emerging evidence suggests that oligodendrocyte precursor cells (OPCs) and oligodendrocytes (OLs) are important contributors to PNN formation and remodeling ^46–53^. However, most of the literature on OL modulation of neural activity has focused on activity-dependent changes in myelination after learning ^54–64^. Here we asked whether oligodendrocyte lineage cells (OLCs) modulate PV IN activity beyond their canoncial role as facilitators of electrical conduction^65–75^.

We address these questions using the model of safety compared to fear learning. We discover that safety conditioning results in long-term reduction of contextual fear, which is associated with the recruitment of OPCs to a satellite position at PV cells in the IL, followed by their maturation and the breakdown of the PNNs surrounding the PV-satellite OL pairs. Furthermore, using optogenetic inhibition, we demonstrate that these changes depend on IL PV activity during safety conditioning. Therefore, we propose that safety learning promotes a program of satellite OL-PV communication that drives PNN degradation, thereby promoting plasticity in the IL.

## Results

### Safety conditioning attenuates remote contextual fear memory

Here we adopted a previously described safety conditioning protocol^76,77^, consisting of mice receiving a shock to the paws in the training context at any time except during the presentation of an auditory stimulus, indicating a period of explicit safety. After habituation to the cue and the training context, reporter *PV-Cre-Ai9* male mice (C57B6J) were exposed to safety conditioning (SC, n=12) and their behavior was compared to contextual fear conditioning alone (CFC, n=10) and to cue-alone exposure (CA, n=10). All mice were first habituated to the context, where the SC and CA groups were also habituated to five presentations of the conditioned stimulus (CS) consisting of a 1-sec house light that co-occurred with a 30 sec tone (4kHz, 50ms pips, occurring at 1Hz for 30s, **Figure 1A-B**, **Supplementary Figure 1**, see Methods). Next, mice underwent two conditioning sessions, with the CFC group exposed to five unconditioned stimulus trials (US, 1 sec, 0.6mA foot shocks, 60-120 sec ITI) per session, the CA group exposed to five CS-alone trials per session, and the SC group was exposed to five unpaired US and CS presentations, such that CS signaled the explicit absence of the shock (**Figure 1A-B, Supplementary Fig. 1**). The next day, all groups underwent a recent retrieval session, and 21 days later, a remote retrieval session (**Figure 1A-B, Supplementary Figure 1**). During both retrieval sessions, all groups underwent context retrieval by virtue of being placed back in the training context, and the SC and CA groups also received five CS trials to probe cue-retrieval. Contextual defensive freezing (defined as full cessation of movement for longer than 1 sec, except for beathing) was assessed in all groups by sampling 30sec time-periods prior to each CS delivery in the CA and SC groups with the CFC group yoked to these same times, **Figure 1C, Supplementary Figure 1**), and CS-evoked freezing was assessed in the CA and SC groups.

**Figure 1.**
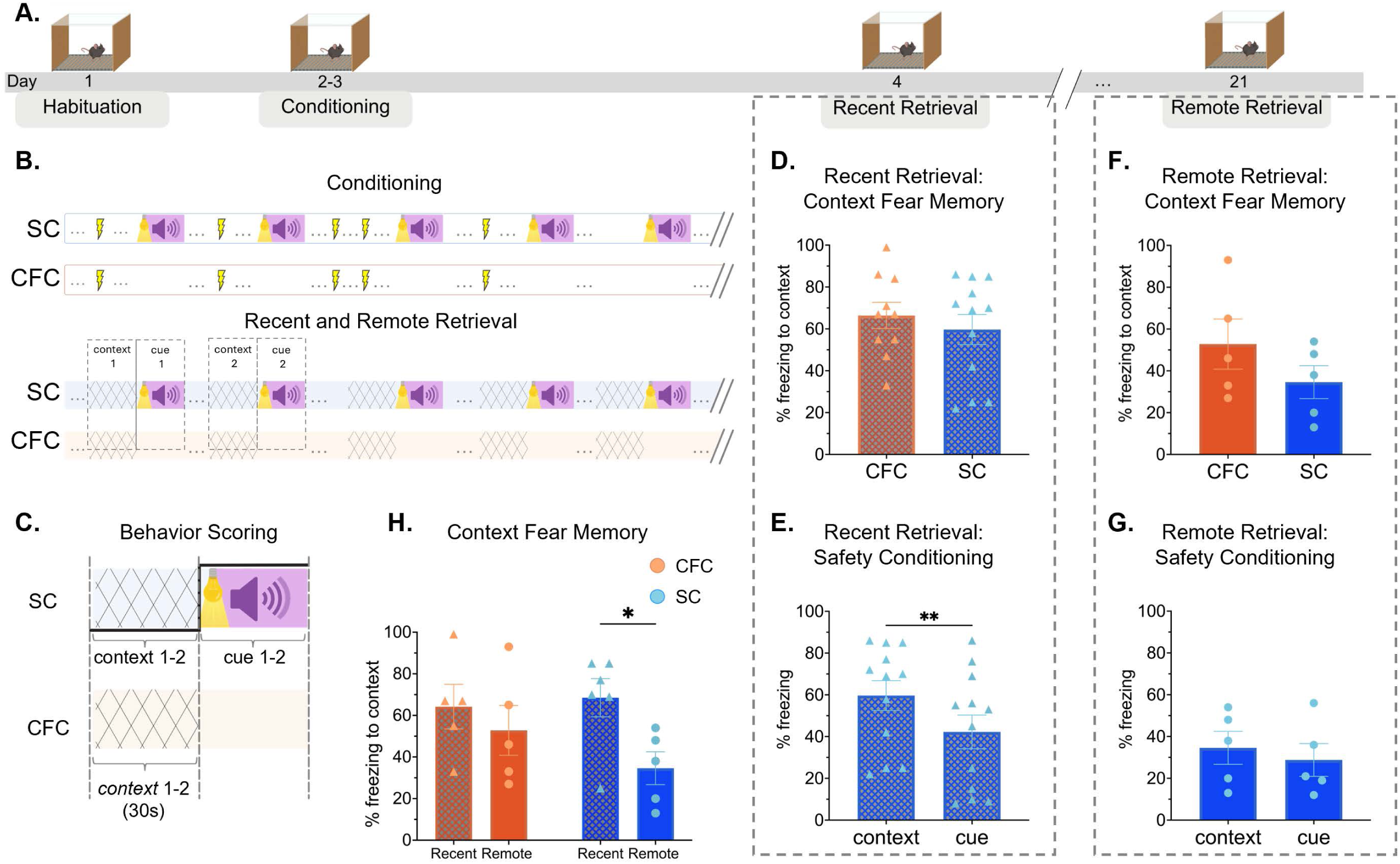
Safety conditioning decreases remote contextual fear memory. **(A)** Behavioral schematic: Day 1: mice were habituated to the training context (CFC, n=10) or to the context and cues (SC, n=12); Day 2-3: conditioning, Day 4: recent retrieval, Day 21: remote retrieval. (**B)** Sample structure of CFC and SC conditioning and retrieval sessions. During each conditioning session, the SC group received five unsignaled foot shocks US (0.6mA, 1 sec) and five CS (1-sec house light co-occurring with 4kHz, 50ms pips playing for 30sec at 1Hz) that were never paired with the US. The CFC group received five unsignaled foot shocks (0.6mA, 1 sec). During retrieval sessions, the SC and CFC groups we re-exposed to the conditioning context with the SC group received five CS trials. **(C)** Schematic showing time bins during which percent freezing was calculated. Percent contextual freezing was assessed in the 30sec prior to CS 1 and 2 of the SC group. The same time bins were identified and assessed for CFC group. Percent freezing during the CS was assessed during the 30 sec cue presentation. **(D)** At recent retrieval, contextual freezing in CFC and SC groups is the same (unpaired t-test, p=0.5). **(E)** At recent retrieval, the CS inhibits contextual freezing in the SC group (paired t-test, p<0.01). **(F)** At remote retrieval, contextual freezing in CFC and SC group is the same (unpaired t-test, p=0.24). **(G)** At remote retrieval, the SC group shows similarly low defensive freezing to the CS and the context (paired t-test, p=0.23). **(H)** Contextual freezing decreases from recent to remote retrieval in the SC but not the CFC group (two-way rm-ANOVA: timepoint, F(1,8)=7.62, p=0.025, Sidak’s multiple comparison, SC group x timepoint, p<0.05). Bars represent mean ± SEM.

During the recent retrieval session, there was no significant difference in contextual fear between the CFC and SC groups (unpaired t-test, p=0.5, **Figure 1D**), and both groups showed more contextual freezing than the CA group (**Supplementary Figure 1**), indicating that for the SC and CFC groups, the shock had rendered the context more aversive than baseline. For the SC group, cue presentation significantly decreased defensive freezing in the training context (paired t-test, p=0.001, **Figure 1E**), whereas cue presentation had no effect on behavior in the CA group (paired t-test, p=0.65, **Supplementary Figure 1**), indicating that for the SC group, there was a conditioned inhibitor of fear.

During remote retrieval, there was no difference in contextual freezing between the SC and CFC groups (**Figure 1F**, unpaired t-test, p=0.24). However, when we compared contextual freezing in the two groups across the recent and remote retrieval timepoints, we identified a statistically significant effect of timepoint (two-way rm-ANOVA, F(1,8)=7.616, p=0.025). A post-hoc comparison revealed that in the SC group, contextual freezing decreased from recent to remote retrieval (Sidak’s, p<0.05, **Figure 1H**), whereas in the CFC group, contextual freezing remained the same across both timepoints (p=0.35, **Figure 1H**). Notably, in the SC group, defensive freezing at remote retrieval was similarly low for context and cue (paired t-test, p=0.24, **Figure 1G**). Thus, at recent retrieval, the SC and CFC groups showed more contextual freezing than the CA group, at remote retrieval, only the CFC group showed significantly more contextual freezing than the CA group (**Supplementary Figure 1D-E**). Given that contextual freezing in the CA group was low at both time points (p=0.4, **Supplementary Figure 1F**), we conclude that the SC group significantly decreased contextual fear from the recent to remote retrieval, suggesting that at remote retrieval, the SC group had updated the memory of context to be less aversive.

### At remote safety retrieval, a greater proportion of IL PVs are active and not surrounded by PNN

Due to the importance of PV activity in shaping cortical activity, gamma oscillation dynamics, and cognitive computations^29,30,78–83^, we focused on PV INs in the IL to investigate the possible cellular mechanisms underlying changes in contextual fear from recent to remote retrieval in the SC group. Frist, we used immunohistochemical staining of brain sections from our *PV-Cre-Ai9* reporter mice with antibodies specific for the immediate early gene cFos at recent and remote retrieval to quantify the proportion of active PV INs (PV:cFos/PV, **Figure 2A)**. A two-way ANOVA revealed no main effect of timepoint (F(1, 9) = 2.906, p=0.12) or group (F(1, 8) = 0.03249, p=0.86), but a timepoint x group interaction (F(1, 8) = 23.97, p=0.001). Multiple comparisons showed that at recent retrieval, the CFC groups showed higher PV activity than the SC group (p=0.01). Further, PV activity signiricantly decreased from recent to remote retrieval in the CFC group (p<0.001), such that at remote retrieval, a significantly larger proportion of PV INs were active in the SC than CFC group (p<0.001, **Figure 2B**).

**Figure 2.**
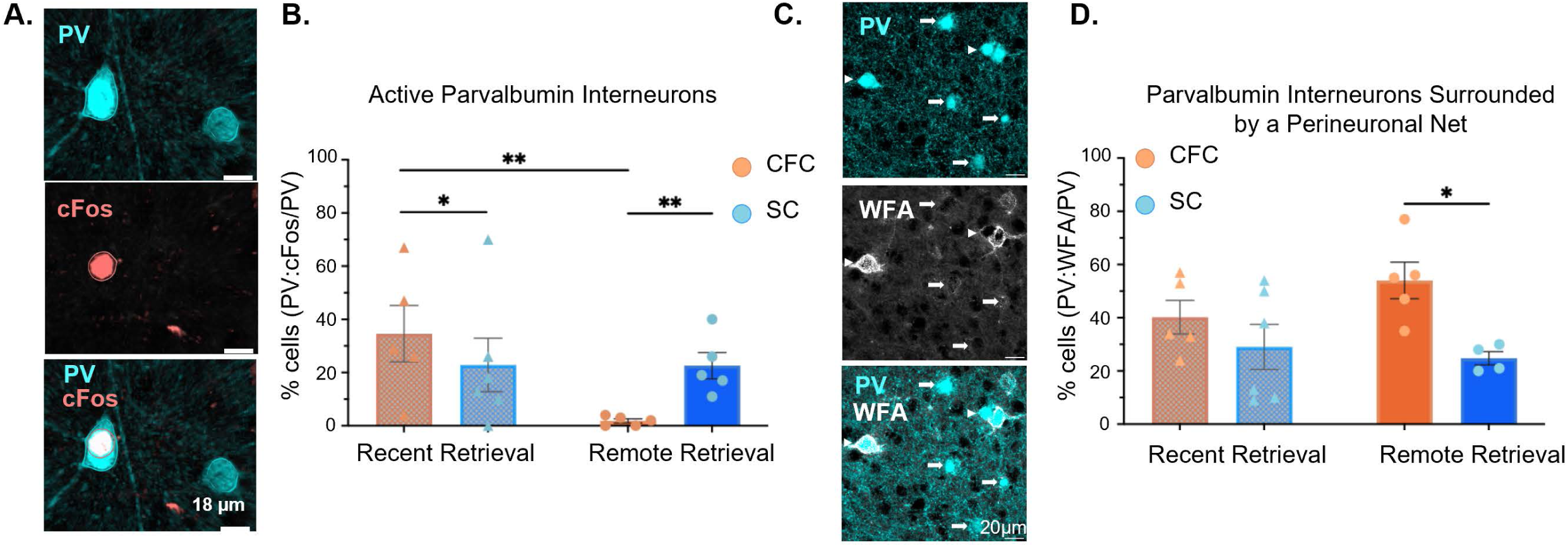
At remote retrieval, the safety conditioned group has higher IL PV activity, and fewer PNNs around PVs than the contextual fear group. **(A)** Example of a PV cells (cyan) expression cFos+ (red). **(B)** At remote retrieval, there is a higher percentage of active PV cells (PV:cFos+/PV) in the IL of the SC than the CFC group, with decreased activity in PV cells of the CFC group from recent to remote retrieval (two-way ANOVA, group x timepoint F(1,8)=23.97, p=0.001, Uncorrected Fisher’s LSD multiple comparisons: Remote retrieval SC vs CFC, p=0.007, CFC recent vs remote, p=0.005, recent retrieval CFC vs SC p=0.01). **(C)** Example of PV cells (cyan) with PNNs labeled by Wisteria floribunda agglutinin (WFA, white). Arrows, PV without PNN; arrowhead, PV with PNN. **(D)** At remote retrieval, a lower percentage of IL PV cells is surrounded by a PNN (PV:WFA+/PV) in the SC than CFC group (two-way ANOVA, group F(1,7)=9.54, p=0.018, Uncorrected Fisher’s LSD multiple comparisons: remote retrieval CFC vs SC p=0.02).

We then asked whether these differences in PV activity were associated with the presence of PNNs around the PV IN. We used Wisteria floribunda agglutinin (WFA), a marker for the N-acetylgalactosamine, to identify and quantify the percentage of PV INs covered by PNN during recent and remote memory retrieval (**Figure 2C**). A two-way rm-ANOVA showed a main effect of group (F(1,7)=9.539, p=0.018), but not of timepoint (F(1, 9) = 0.3365, p=0.57), or group x timepoint interaction (F(1, 7) = 1.760, p=0.22). Multiple comparisons indicated that at remote retrieval, the safety group was characterized by a significantly smaller percentage of PV IN that were covered by PNN (PV:WFA/PV) compared to the CFC group (p= 0.02, **Figure 2D**).

To gain better insight into the relationship between the presence of PNNs and PV activity, we quantified the percentage of active PV INs surrounded by PNNs (PV:WFA:cFos/PV:WFA) and found that PV cells surrounded by PNNs were predominantly inactive, regardless of behavioral group or timepoint **(Supplementary Figure 2**). This result is consistent with previous reports on the role of PNN in constraining PV activity^84–86^. We next directly compared the activity profile of the PV population with versus without PNNs at remote retrieval. Given that in the CFC group, all sampled PV cells were covered by PNNs whereas the SC group showed variability in PNN coverage (data not shown), we evaluated this question only in SC mice (n=5). This analysis showed that in the PV population without PNNs, a significantly greater proportion of cell was cFos+ (PV:WFA-:cFos+ vs PV:WFA:cFos-, Chi Square, p<0.05; **Supplementary Figure 2**). However, despite PV cells without PNNs having a higher percentage of active cells, inactive PV cells (PV:WFA-:cFos-/PV:WFA) were still the majority, t-test, p<0.05; **Supplementary Figure 2**).

Overall, at remote retrieval, the SC group showed fewer PV cells surrounded by PNNs, and a more varied pattern of activity than the CFC group, due to PVs without PNNs having a more variable activity profile. These findings suggest that in the SC group, at remote retrieval, a greater proportion of IL PVs lack PNNs and are in a higher state of plasticity, likely able to respond to presynaptic inputs that can up- or down-regulate their activity. Conversely in the CFC group, the higher proportion of PVs surrounded by PNN was consistent with inactivity and less plasticity.

### Safety learning recruits oligodendrocytes without increasing myelination in the IL

To explore the potential molecular mechanisms underlying behavioral changes detected after SC, we analyzed the mPFC transcriptome in a separate cohort of safety and cue-fear conditioned mice at recent and remote time points (see Methods, **Supplementary Figure 3**). We observed a significant upregulation of myelin- and oligodendrocyte-transcripts in the mPFC of safety-compared to fear-conditioned mice. Among these, the SC group showed increased expression of *Enpp6*, a gene associated with newly formed OLs (NFOLs) at the recent (**Supplementary Figure 3A-C**) and remote timepoints (**Supplementary Figure 3D-I**), whose higher expression levels in the safety group was also validated using qPCR and RNAScope (**Supplementary Figure 3**). Notably, we also observed higher levels of transcription factor *Myrf*, and myelin genes *Mbp* and *Mobp* in the safety relative to fear group at the remote timepoint (**Supplementary Figure 3E-I**), and therefore asked whether there were differences in the overall myelination of the mPFC in the SC compared to the CFC group at either recent or remote retrieval, but found that there were no differences (**Supplemental Figure 4**). Thus, the experience-dependent increase in OL transcripts detected in the safety group relative to the fear group, was not linked to an experience-dependent increase in myelination, and suggested an alternative, non-myelinating function for OLs in safety learning.

### In the IL, more satellite oligodendrocytes pair with PV cells that lack PNNs after safety conditioning than after contextual fear conditioning

Given the transcriptional changes in OL-specific genes detected after safety relative to cued fear training, we used immunohistochemistry and antibodies for the pan-lineage marker OLIG2 to identify all the cells in this lineage (**Figure 3A**) to inquire whether there were any training related changes in the overall density of OLIG2+ cells after SC vs CFC at either timepoint. This analysis did not detect any difference in the overall density of OLIG2*+* cells in the IL at either recent or remote retrieval between the two groups (**Figure 3B**).

**Figure 3.**
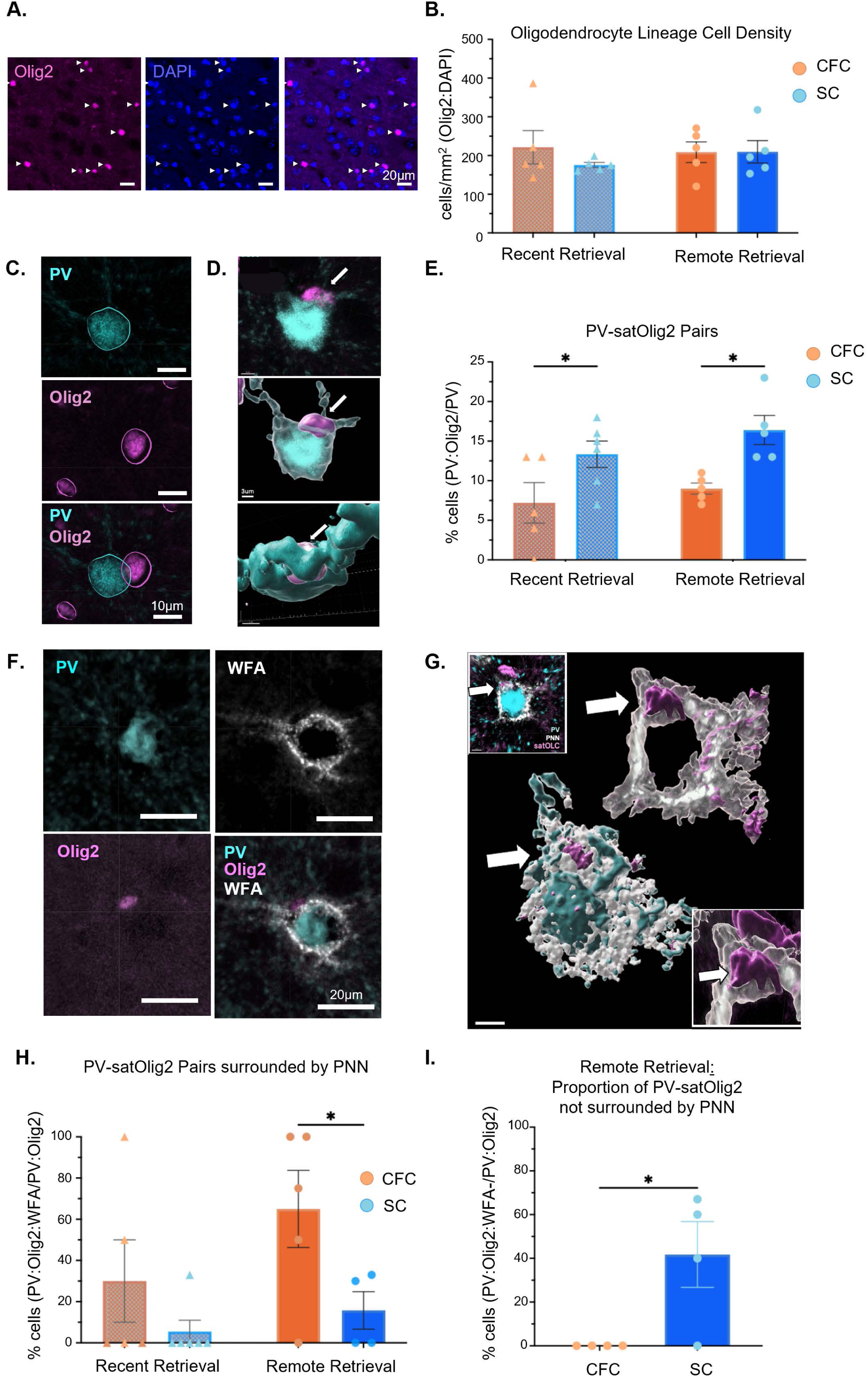
Safety conditioning increases satellite oligodendrocytes pairing with PV INs that do not have PNNs. **(A)** Example of oligodendrocyte lineage cells (OLC) staining with Olig2 (magenta) overlapping with the nuclear DAPI stain (blue) in the IL. **(B)** IL OLC density (Olig2+:DAPI/mm^2^) is the same in CFC and SC groups at recent and remote retrieval (two-way ANOVA, group x timepoint F(1,8)=0.703, p=0.43). **(C)** Example of a PV IN (cyan) with a satellite Olig2*+* (satOlig2+) cell (magenta). **(D)** IMARIS 3D rendering of a PV-satOlig2+ pairing (arrow points to the satOlig2+ cell). **(E)** A higher percent of PV cells are paired with a satOlig2+ (PV:Olig2+/PV) in the SC than CFC group at recent and remote retrieval (two-way ANOVA, group F(1,17)=13.9, p=0.0017, Uncorrected Fisher’s LSD multiple comparisons: recent retrieval, CFC vs. SC p=0.026, remote retrieval CFC vs. SC p=0.011). **(F)** Immunostaining example and **(G)** IMARIS 3D rendering of a PV-satOlig2+ pair surrounded by a PNN (PV:Olig2*+*:WFA+). Arrows, satOlig2+ cell. **(H)** At remote retrieval, the percent of PV-satOlig2+ pairs surrounded by a PNN (PV:Olig2+:WFA+/PV:Olig2+) is lower in SC than CFC (two-way ANOVA, group F(1,16)=16.30, p=0.023, Uncorrected Fisher’s LSD multiple comparisons : remote retrieval CFC vs SC p=0.038). **(I)** At remote retrieval, the SC group has a higher percentage of PV-satOlig2+ pairs that are not covered with a PNN (PV:Olig2+:WFA-/PV:Olig2+, unpaired t-test p=0.032).

In our analysis we noted that many of the OLIG2+ cells were not randomly distributed, but rather in close contact with PV INs, frequently embedded within a pocket of the PV soma **(Figure 3C-D).** Based on their position, often within the same extracellular space as the PNN, we identified them as satellite oligodendrocyte lineage cell (satOLIG2+ cells), making them well-positioned to directly influence PNN structure and PV INs. Surprisingly, more satOLIG2+*-*PV pairings were detected in the SC than in the CFC group, both at the recent and remote retrieval **(Figure 3E)**. This suggested that satOLIG2+cells were recruited to PV INs soon after safety learning. Given the position of these satOLIG2+cells close to the PNN of the PV IN **(Figure 3F-G**), we also quantified the percentage of PV-satOLIG2+ cell pairs surrounded by PNN. We noticed that at recent retrieval, although group averages were not different, in the SC group, PV-satOLIG2+cell pairs were predominantly not covered by PNNs whereas in the CFC group the percentage was highly variable (F-test of variance, F(11.02), p=0.02, **Figure 3H)**. In contrast, at remote retrieval, the proportion of PV-satOLIG2+cells surrounded by PNNs in the SC group was significantly lower than in the CFC group **(Figure 3H)**, with similar variance in both group (F-test of variance, (F(5.27), p=0.2). In other words, the safety group was characterized by a greater proportion of PV INs that were paired with satOLIG2+ and lacked PNNs **(Figure 3I)**. Thus, at remote retrieval, more PV INs that were associated with satOLIG2+ cells lacked PNNs in the SC group, whereas in the CFC group, more PV INs that were associated with satOLIG2+cells were covered by PNNs.

These findings indicated that in the IL, satOLIG2+ cells are more readily recruited to PV INs after SC, with such pairings stably remaining for weeks post-learning. Furthermore, the lower percentage of PV-satOLIG2+ cell pairs surrounded by PNN in the SC group, suggested that the OLIG2 cells in the two groups may be differentially affecting PNN structure, depending on the behavioral group.

### PNN degradation genes are expressed in mature OL

Oligodendroglia regulate neural function in multiple ways, including metabolic support, regulation of extracellular ion concentrations, neurotransmitter and trophic factor release ^87^. Importantly, oligodendrocyte lineage cells are characterized by maturation-stage specific patterns of gene expression that affect their function, and recent studies suggest that at different stages of maturation, OLIG2+ cells differentially express genes involved in PNN remodeling ^46–52,88^ . Given increased satOLIG2+ pairing with PV cells and the decrease in PNN around them after safety learning, we used open-source scRNAseq data (ABC Brain Cell Atlas, Allen Institute, RRID:SCR_024440) to investigate PNN-related gene expression in oligodendrocyte lineage cells at different stages of maturation (**Figure 4A-D**). We found that, at the progenitor stage, 80% of OPCs in the mPFC express genes related to the formation and assembly of the PNN, such as chondroitin sulfate proteoglycans from the lectican family (e.g. versican (*Vcan*), brevican (*Bcan*), aggrecan (*Acan*)), and other supporting proteins such as tenascin-R (*Tnr*) (**Figure 4A**). We also found that NFOLs (marked by *Enpp6*+) are unique in that they co-express PNN degradation enzymes and PNN structural genes. In contrast, mature OLs (identified by the expression of *Opalin*), predominantly express enzymes responsible for PNN degradation^50,51^, such as the proteases of the *Adamts* family (e.g. *Adamts2*, and *Adamts9,* **Figure 4A-D**).

**Figure 4.**
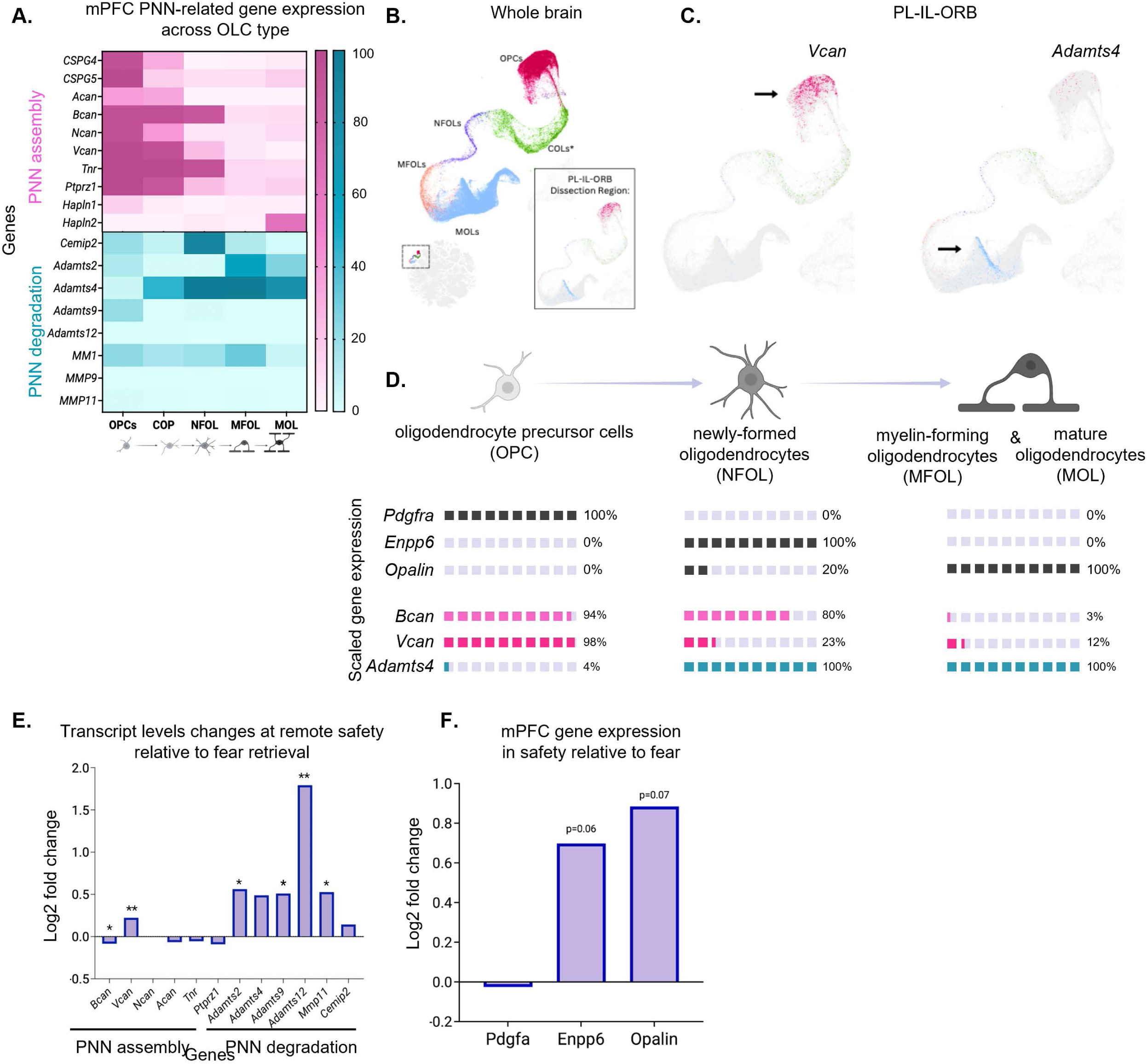
PNN degradation genes are expressed more in mature OLC and are elevated after safety learning. **(A)** Heatmap of differential PNN-related gene expression across the OL lineage; data mined from 10x single-cell RNA sequencing data (Allen Institute’s ABC Atlas; PL-IL-ORB region, male mice), showing the OPC-Oligo Class (#31) subdivided into OPC (#326) and OL (#327) subclasses. The x-axis represents oligodendrocyte lineage stages, including OPCs, committed oligodendrocyte precursors (COPs), NFOLs, MFOLs, and MOLs. The y-axis lists PNN assembly genes (teal) and PNN degradation genes (pink). Brighter colors indicate higher expression levels, with 100% representing full expression within a given cell type. PNN assembly genes are primarily expressed by OPCs and early-stage OLs, whereas PNN degradation genes are expressed by later-stage OLs. **(B)** U-Map of whole-brain scRNAseq gene expression across subdivided OPC-Oligo Classes, with colors distinguishing supertypes: OPCs (red and brown), COPs (green, data not shown again*), NFOLs (purple), MFOLs (orange), and MOLs (light blue). Bottom left shows the OPC-Oligo Class section location in the entire U-Map. Bottom right shows OPC-Oligo overall gene expression in the mPFC (PL–IL–ORB) dissection region. **(C)** Gene expression of PNN assembly gene *Vcan* and PNN degradation enzyme *Adamts4* in OPC-OL class within the PL-IL-ORB region. Arrows indicate the region of highest expression. **(D)** Summary of selected PNN-related gene expression across OPCs, NFOLs, MFOLs, and MOLs in the PL-IL-ORB region (data sourced from the Allen ABC Atlas). Each square represents 10% of gene expression across cell type. Percentages were calculated by importing ABC Atlas UMAP images into ImageJ, measuring expression density (cells/mm²) within each oligodendrocyte supertype, and normalizing to the total density of that supertype. Rows 1-3 are OLC-related genes (grey). Rows 4 & 5 are PNN structural genes *Bcan* & *Vcan* (pink), Row 6 is PNN degradation gene, *Adamts4* (teal). **(E)** Results from RNASeq showing transcription of genes involved in PNN assembly and degradation in the mPFC, two-weeks after safety or cued fear conditioning. A positive log^2^ fold change indicates higher gene expression in safety relative to fear, while a negative value indicates lower expression in SC. Among PNN assembly genes, only versican (*Vcan*) showed upregulation in SC (p=0.009), whereas brevican (*Bcan*) tended to be reduced in safety (p=0.08). Several PNN-degrading enzymes were significantly upregulated in the mPFC after SC: *Adamts2* (p=0.03), *Adamts9* (p=0.04), *Adamts12* (p=0.017), and *MMP11* (p=0.04). Enzymes *Adamts4* (p=0.11) and *Cemip2* (p=0.106) showed a trend towards enhanced expression in the mPFC of SC, but did not reach statistical significance. Asterisks (*) indicate genes with unadjusted p<0.05. **(F)** RNASeq analysis showing transcription levels of oligodendrocyte-lineage genes in safety relative to fear conditioned animals two-weeks after training: *Pdgfrα* is expressed in OPCs, *Enpp6* is expressed in NFOLs, *Opalin* is expressed in myelin-forming and mature oligodendrocytes (MFOLs and MOLs).

This suggests that PNNs may be differentially regulated by OLCs, depending on their maturation status, which shifts with experience. Turning to our bulk RNASeq data in the mPFC, we analyzed OLC and PNN-related gene transcription in remote safety vs cued-fear conditioned mice. This analysis revealed that in the mPFC of safety conditioned mice, transcription of PNN degradation enzymes was significantly upregulated and transcription of PNN assembly genes (e.g., lecticans) was significantly downregulated (**Figure 4E**) relative to cued-fear conditioned mice. Likewise, we saw higher transcription of newly formed (*Enpp6+*) and mature (*Opalin+*) OLs in safety relative to cued-fear conditioned mice (**Figure 4F**). These findings suggest that the maturation status of satOLIG2+ cells could alter the status of PNNs surrounding PV cells in the IL, with more mature satOLIG2+ cells signaling PNN breakdown after SC than CFC.

### PV INs consistently pair more with mature satellite OLs at remote retrieval only after safety learning

To track satellite OLC maturation status in the mPFC at different timepoints after SC and CFC learning, we used immunohistochemistry and antibodies for the pan-lineage marker OLIG2 to identify all the cells in this lineage, NG2 to identify OPCs and CC1 to identify mature OLs **(Figure 5A)** at recent and remote retrieval. We conducted a detailed analysis of the maturation status of the PV-paired satOLIG2+ cells (**Figure 5B-C**) and found that at recent retrieval, the cumulative density of satOPCs (Olig2+:NG2+:DAPI+) and satOLs (Olig2+:CC1+:DAPI+) didn’t differ in the CFC (Kolmogorov-Smirnov (K-S) test, p=0.46) or the SC (p=0.57) groups (**Figure 5D-E**), such that on average their numbers were the same in both groups (**Figure 5F-G**). At remote retrieval, the cumulative density of satOPCs and satOLs was also similar in the CFC group (K-S test, p=0.98, **Figure 5H**), whereas in the SC group, the cumulative density of satOLs was significantly right-shifted (K-S test, p=0.016) from the satOPCs, indicating that satOLs constituted a larger proportion of the satOLIG2+ cells (**Figure 5I**), such that on average there were more satOLs than satOPCs (Mann-Whitney, p=0.039) in the SC group but no differences in the percent of mature vs immature satOLIG2+ cells in the CFC group (**Figure 5J-K)**. Interestingly, when we compared changes between recent and remote retrieval in the SC group, we found that the percent of mature satOLIG2+ cells (CC1+) didn’t change, but the proportion of OPCs showed a trend towards a decrease (Mann-Whitney, p=0.07).

**Figure 5.**
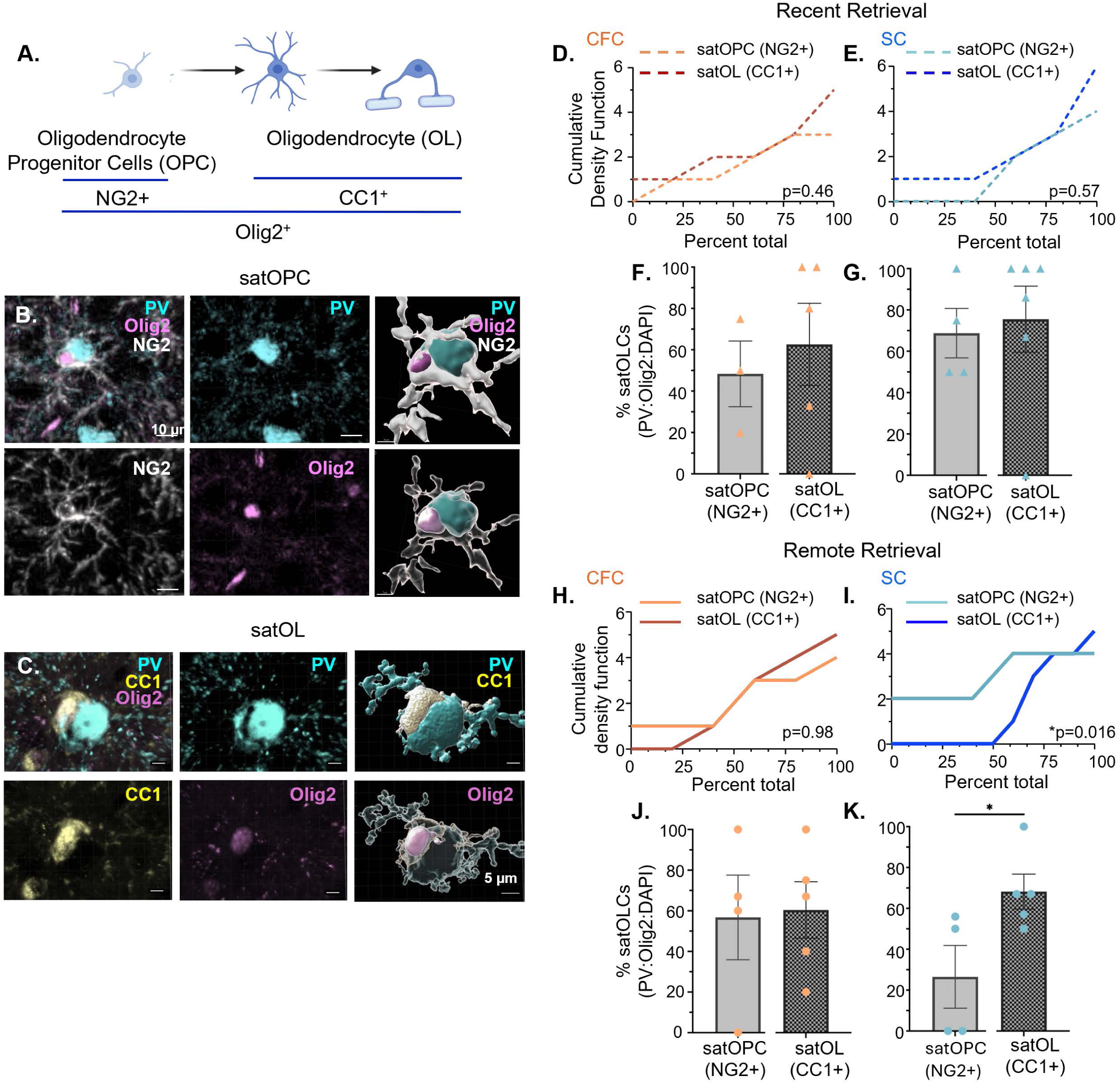
At remote retrieval, PV INs pair more with mature satellite OLs after safety learning. **(A)** Schematic of OLC markers at different maturation stages. Olig2 is a pan-lineage marker, NG2 identifies oligodendrocyte progenitor cells (OPCs) and CC1 identifies mature OLs. **(B)** satOPC labeling: Immunolabeled example (*left*) and IMARIS 3D rendering (*right*) of an OPC (NG2+, white) satellite oligodendrocyte (Olig2+, magenta) (satOLIG2+) in tight proximity of its host PV cell (cyan). **(C)** satOL labeling: Immunolabeled example (*left*) and IMARIS 3D rendering (*right*) of a mature (CC1+, yellow) satellite oligodendrocyte (Olig2+, magenta) (satOLIG2+) in tight proximity of its host PV cell (cyan). **(D-G)** satOLIG2+ maturation status at recent retrieval. (**D**) At recent retrieval, the cumulative density distribution of satOPCs and satOLs were similar in the CFC group (p=0.46) and the (**E**) SC group (p=0.57). (**F**) Similar average proportions of satOPCs (light grey) and satOL (dark grey) paired with PV INs in the CFC and (**G**) SC groups at recent retrieval. (**H-K**) satOLIG2+ maturation status at remote retrieval. (**H-I**) At remote retrieval, the cumulative density distribution of satOPCs and satOLs was similar in the CFC group (p=0.98), but in the SC group (**I**) the satOL distribution was significantly right-shifted (p=0.016) from the satOPC distribution, indicating that satOLs comprise a larger proportion of the population. (J) Similar average proportions of satOPC (light grey) and satOL (CC1+, dark grey) paired with PV INs in the CFC group but (**K**) more satOLs than satOPCs (Mann-Whitney, p=0.039) paired with PV INs in the SC group at remote retrieval.

These findings suggest learning-dependent patterns of satOLIG2+ maturation in the IL, whereby satOLIG2+ cells are recruited to a satellite position on PV INs in various stages of maturation after both types of learning, likely expressing PNN assembly and degradation genes. However, as time passes, the satOLIG2+ cells that are found in the CFC group continue to be in a variety of maturation status, whereas, in the SC group, the satOPCs decrease in number, leaving the majority of satOLIG2+ cells in a mature state (**Figure 5G**), likely expressing genes that facilitate PNN breakdown (**Figure 4)**.

### PV activity during safety learning is necessary for recruitment and maturation of satellite oligodendrocytes, PNN remodeling and decreased contextual fear at remote retrieval

PV-mediated signaling via γ-amino-butyric acid (GABA) was previously shown to recruit OLCs and spur myelination^70,89^. Thus, to determine whether PV activity contributes to satOLIG2+ recruitment in safety learning, we used optogenetics to inactivate PV INs in the IL. We injected the Cre-dependent inhibitory opsin *enhanced Archeorhodopsin 3.0* (eArch, n=8) or its control virus, eYFP (n=6), in the IL of male mice (F1 of *PV-Cre* C57B6J and wild-type 129SvEv mice). Three weeks after viral injection, mice were handled and habituated to the training context, the tether, light (on/off ramp-modulated light delivery^91^), and the safety cue, and then underwent safety conditioning with PV IN inhibition only during the safety cue (**Figure 7A-B**). PV IN inhibition did not affect the ability to learn safety, and during recent retrieval eYFP and eArch groups showed significantly reduced freezing during the safety cue compared to context alone (**Figure 7C**). Note that F1 C57B6J/129SvEv mice typically show higher defensive freezing than wild type C57B6J mice shown in figure 1, but both strains decrease defensive freezing during the safety cue relative to context. Indeed, inhibiting PV INs during learning resulted in a more robust decrease in freezing during the safety cue at recent retrieval. At remote retrieval, the eYFP group no longer showed suppressed freezing during the safety cue relative to context alone (**Supplementary Figure 5**, as observed earlier in the wild type (**Figure 1G**). However, the eArch group continued to show suppressed freezing during the safety cue relative to context (**Supplementary Figure 5**). This difference was attributable to decreased contextual freezing from recent to remote retrieval in the eYFP group but continually high contextual freezing at both the recent and remote retrieval timepoints in the eArch group (**Figure 7D**). This suggested that IL PV activity is not necessary for acquiring the safety cue association, but plays a critical role in decreasing contextual freezing that emerges at remote retrieval after safety conditioning.

**Figure 6.**
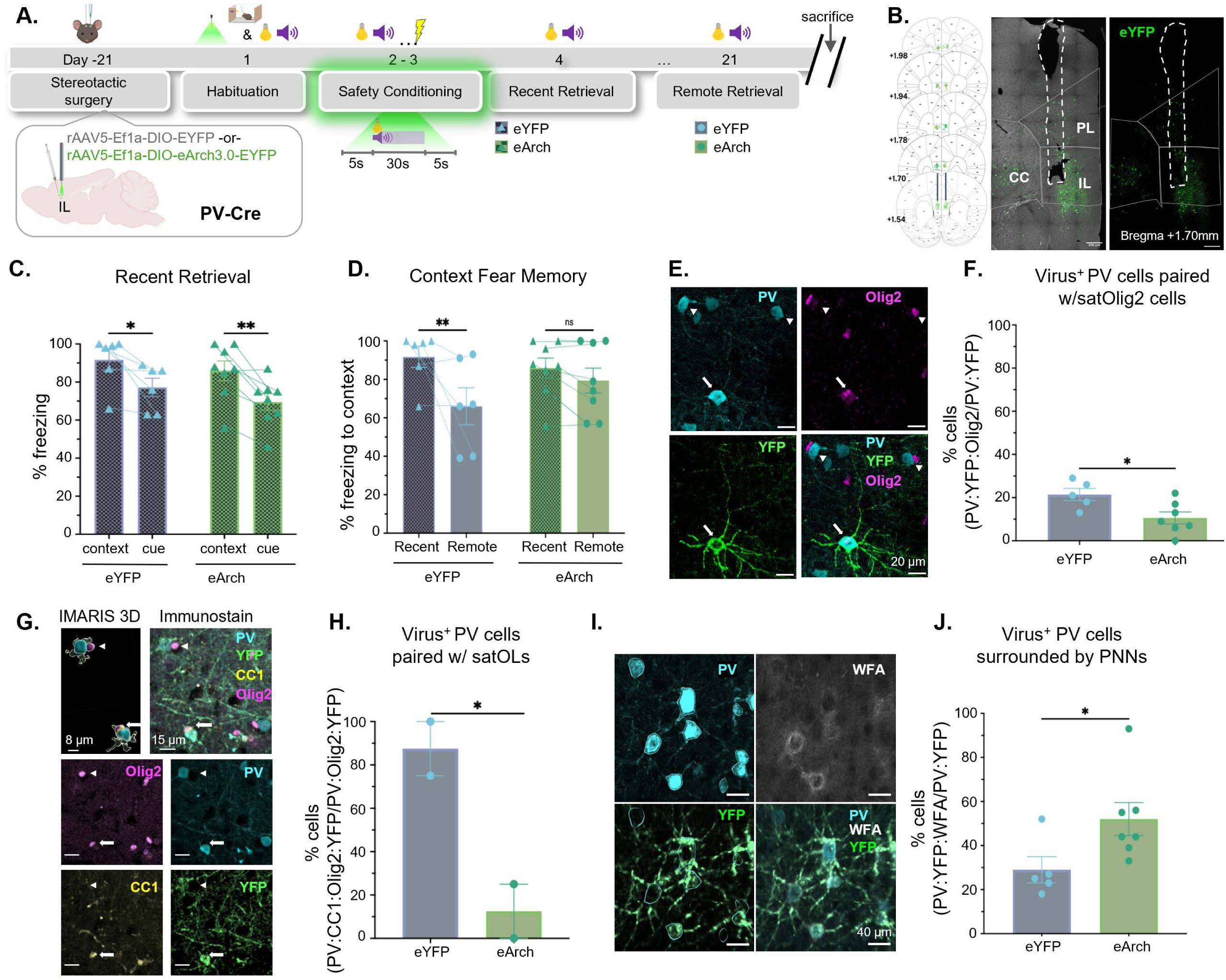
PV IN activity during safety learning is necessary for satellite OL recruitment and maturation, PNN degradation and decreased contextual fear at remote retrieval. **(A)** Experimental timeline: In male PV-Cre mice, 0.3 µl of the inhibitory virus r_AAV5_-EF1a-DIO-eArch3.0-EYFP (eArch) or its control (r_AAV5_-Ef1a-DIO-EYFP, eYFP) was injected in the IL, and optic fibers were implanted in the IL. Following 21 days of recovery and viral expression, mice were habituated to context, cue and laser and then received two days of safety conditioning with the laser on during the safety cue. The next day, mice received a recent retrieval session, and 21 days later, a remote retrieval session. Mice were perfused after the remote retrieval session for verification of targeting and immunohistochemistry. (B) *Left,* Mapping of virus placement in the IL for all subjects. *Right*, Example of viral injection (eYFP, green) and fiberoptic placement in the IL. **(C)** Percent freezing is lower during the safety cue than context alone in the eYFP and eArch groups (two-way ANOVA, trial type, F(1,12)=24.1, p=0.0004, group F(1, 12)=1.10, p=0.31, trial type x group F(1,12)=0.08, p=0.77; Uncorrected Fisher’s LSD multiple comparisons: eYFP context vs safety cue p=0.01, eArch context vs safety cue, p=0.002). **(D)** Percent freezing to context alone decreases from recent to remote retrieval in the eYFP but not in the eArch group (two-way ANOVA, timepoint F(1,12)=8.77, p=0.019; group F(1,12)=0.23, p=0.64; timepoint x group, F(1,12)=3.11, p=0.103; Uncorrected Fisher’s LSD multiple comparisons, eYFP recent vs remote, p=0.008, eArch recent vs remote, p=0.38). **(E)** Example of a virus-expressing PV cell paired with a satOLIG2+ cell (PV+:eYFP+:Olig2+). Arrowheads, PV cells (cyan) paired with satOLIG2+ cells (magenta), and no viral expression, Arrow, PV cell with viral expression (eYFP, green) that is paired with a satOLIG2+ cell. **(F)** At remote retrieval, the percentage of eYFP-expressing PV cells that are paired with satOLIG2+ cells is higher in the eYFP than the eArch group (unpaired t-test, p=0.02). **(G)** (*Top* Left) IMARIS 3D rending and Immunohistochemistry staining showing an example of a PV cell (YFP+) paired with a CC1+ satOLIG2+ cell (arrow, PV:YFP+:Olig2+:CC1+), and (arrowhead) shows an example of a PV cell (YFP+) paired with a CC1-satOLIG2+ cell. **(H)** Percent of mature satOLIG2+ cells paired with eYFP+ PVs is higher in the eYFP than the eArch group (eYFP+:PV:Olig2+:CC1+/eYFP+:PV:Olig2+, unpaired t-test p=0.05). **(I)** Example of eYFP+ PV cells surrounded by a PNN (PV:eYFP+:WFA+). **(J)** A higher percentage of eYFP+ PV cells are surrounded by a PNN (eYFP+:PV:WFA+/eYFP+:PV) in the eArch than the eYFP group is (unpaired t-test, p=0.048). All data is plotted as means ± SEM.

**Figure 7.**
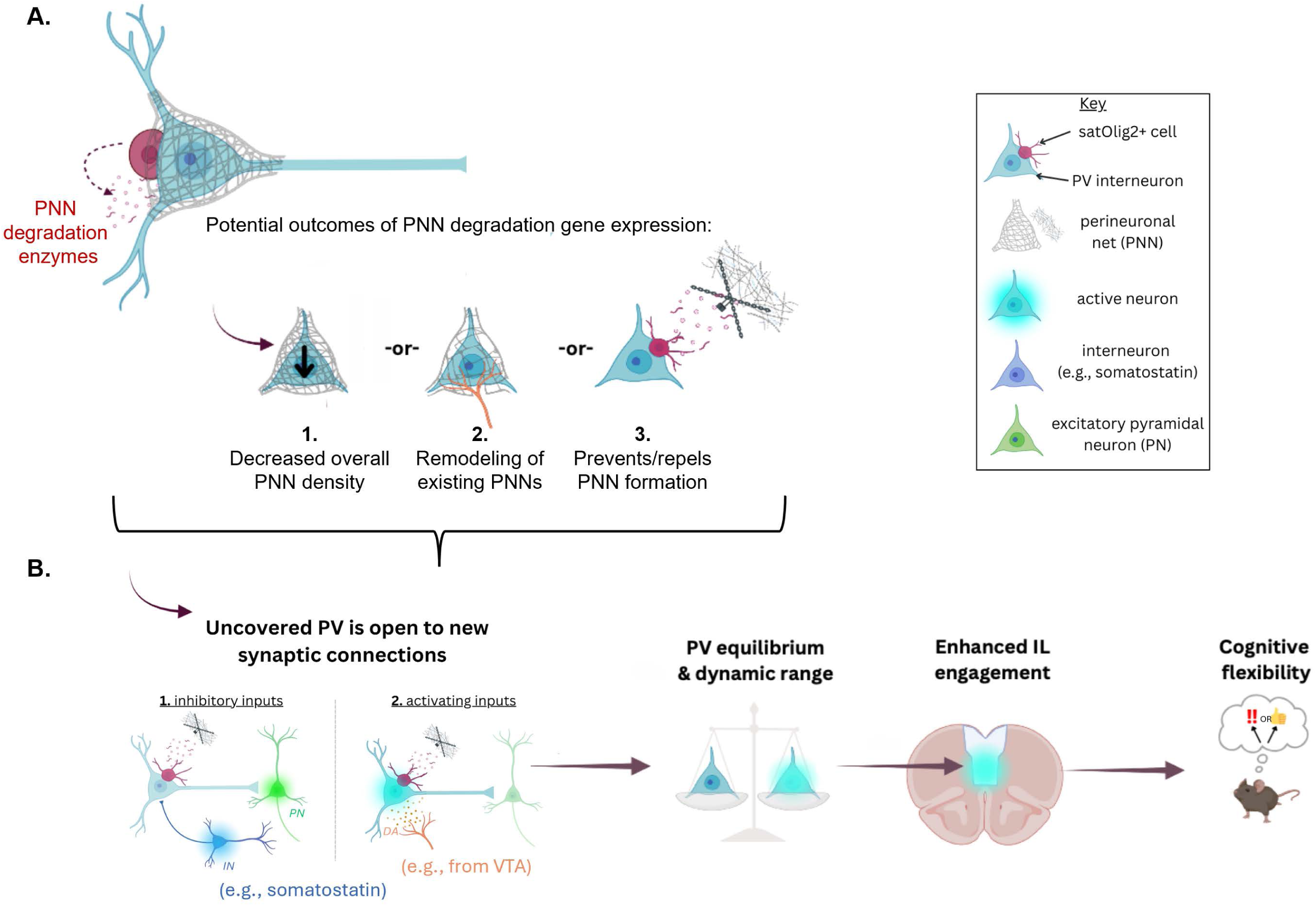
Model of experience-dependent satOLIG2+-PNN regulation in the IL. **(A)** Safety conditioning triggers mature satOLIG2+ cells to transcribe genes for PNN degradation enzymes. This activity could have a range of effects, such as (1) decreasing overall PNN density, (2) remodeling existing PNNs to increase holes, creating more space for new synaptic input or decreasing existing inputs, or (3) preventing PNN formation **(B)**. With reduced PNN presence, PV INs are open to new synaptic connections, both activating and inhibitory (e.g. as in Wang et al., 2024^33^, from somatostatin-expressing interneurons). This way, PV activity is kept at an equilibrium, allowing for IL activity during fear suppression. By limiting PNN expression, PV INs gain an expanded dynamic range, allowing them to selectively fire, based on behavioral demands, fostering adaptive learning and cognitive flexibility.

To investigate whether PV activity during safety learning affected satOLIG2+ cell recruitment to the PV INs, we quantified the percentage of virus-expressing PV INs that were paired with satOLIG2+ cells at remote retrieval (**Figure 7E**). This analysis showed that despite similar levels of viral expression in both groups (**Supplementary Figure 5**) inhibiting PV activity during learning reduced satOLIG2+ cell pairing with PV INs in eArch mice relative to the eYFP controls (**Figure 7F**), indicating that PV activity at learning recruits satOLIG2+ cells to PV INs. We then evaluated the maturity status of satOLIG2+ cells paired with PV INs in both groups. This comparison showed that in the eYFP control group, there were more mature satOLIG2+ cells associated with PV INs than in the eArch group (**Figure 7G-H**), indicating the PV IN activity during safety learning also facilitated satOLIG2+ cells maturation into OL. Finally, we evaluated whether PV activity affected PNN expression in both groups and found that there were fewer PV INs surrounded by PNNs in the eYFP than in the eArch group (**Figure 7I-J**), indicating that PV activity during safety learning contributes to decreased PNN expression at remote retrieval. Taken together, these findings show that PV activity in the IL during safety learning is critical for the recruitment and maturation of satOLIG2+ cells, and the long-term degradation of PNNs around PV INs, aiding behavioral flexibility.

## Discussion

This study identifies a neuroglial mechanism that differentially regulates PV interneuron activity in the IL of the mPFC during safety and fear learning. Our data suggest activity-dependent recruitment of oligodendrocyte lineage cells to the soma of active PV interneurons, where, depending on their maturation status, they may favor the composition or degradation of the PNN. Whereas safety learning is associated with a greater proportion of mature OLs soon after learning, and a smaller percentage of PV IN surrounded by PNN at remote retrieval, fear learning is characterized by the persistence of PNN surrounding PV cells. Since the absence of PNN renders interneurons more responsive to presynaptic inputs, we conclude that safety learning is characterized by greater PV IN plasticity, whereas fear learning is characterized by less PV plasticity, consistent with the behavioral rigidity of fear memories. These findings offer new insights into a non-myelinating role for oligodendrocytes in updating long-term fear memory and the dynamic relationship between PV, PNNs, and OLs.

### Role of Satellite Oligodendrocytes in PNN Plasticity

We demonstrate that SC leads to recruitment of OLIG2 cells to PV IN in the IL, which is driven - at least in part - by IL PV activity during learning and leads to PNN remodeling. It has been previously shown that OPC express mRNA for PNN structural components, while mature OLs are enriched in the expression of PNN degradation enzymes^46–52,88^. The NFOL characterized by *Enpp6* expression, in contrast, show a unique transcriptome, characterized by the co-expression of both PNN structural and PNN degradation genes (ABC Atlas, Allen Institute). Our data show that the differentiation of OPC into OL during safety learning was dependent on PV activity, consistent with previous reports ^90^. However, the fact that the higher levels of *Enpp6* and myelin genes in the safety group, were not paralleled by global changes in myelination in the IL region of the PFC, also suggested a non-myelinating role for these newly formed OL.

The detection of enhanced oligodendrogenesis shortly after learning, is consistent with previous reports on other forms of learning^60,61^. These data were previously discussed in terms of adaptive myelination in response to learning and indeed both SC and CFC groups were characterized by similar levels of myelin in the IL at remote retrieval. Our analysis extends previous findings, as we report that only the safety learning group showed higher percentages of CC1+ mature OL at recent recall, likely resulting from the progression of *Enpp6+* NFOLs and later contributing to PNN remodeling around PV cells.

We suggest that this remodeling process is highly localized, as OLIG2+ cells can be found in close apposition to the soma of PV in the IL. These satellite OLIG2*^+^* cells may influence PNN formation or degradation, depending on their maturation status, because they make direct contact with neurons and often occupy the same perineuronal space as the PNNs. Originally described by del Río Hortega in 1928 ^93^, satellite oligodendroglial cells have long been described and characterized morphologically^97^ and cytologically ^98,99^, although their function remains elusive. Limited evidence suggests that satellite oligodendroglia may support their host neurons by maintaining homeostasis^98,100^ and directly regulate membrane potential during high frequency firing^96,101^, which suggests their potential function in fine-tuning PV activity and information processing in the IL. Moreover, post-mortem studies have linked a reduction in prefrontal satOL density to psychiatric disorders such as schizophrenia^102–105^, highlighting a potential role for these cells in cognitive processing. Our findings suggest a learning- and memory-dependent role for satellite OLIG2*^+^*cells, underscoring their pivotal impact on shaping behavior.

Importantly, we found that satellite OLIG2*^+^* cells are more readily recruited to PVs following SC, when animals are less fearful relative to the CFC group. The PV-satellite OLIG2*^+^* cell pairing established soon after safety learning, remained upregulated for weeks and were rarely characterized by the presence of PNNs covering PV-paired with satellite OLIG2*^+^* cells during remote safety retrieval. In contrast, CFC animals displayed higher fear and had fewer PV-satOLIG2+ pairs at both retrieval timepoints, with most PV-paired with satOLIG2*^+^* cells being entirely encased in PNNs at remote retrieval. More PNNs in this group corresponds to restricted PV activity, which may be responsible for preventing the neural plasticity necessary for updated fear responses, contributing to the enhanced contextual fear memory retention seen in the CFC group.

Notably, mature OLs express genes for PNN-degrading enzymes (i.e., *Adamts* family), and in the SC group, we observed a prevalence of mature satOLs relative to the OLC population. Thus, we propose that SC promotes the expression of these genes by satOLs to enable PNN degradation and enhance synaptic plasticity early in memory formation. The recruited OLs, likely NFOLs, express both PNN components and PNN-degrading enzymes. We hypothesize that NFOLs are potentially in an OPC-like state during recruitment to active PV cells and rapidly differentiate near PV cells involved in the memory trace. Because our inhibition data suggest that satOLIG2+ cells are primarily recruited to PVs that are active during learning, they are likely not pairing with INs that are already covered with a PNN, which are inherently less active. Thus, instead of degrading existing PNNs, satOLs likely prevent PNN formation around a subset of PV cells, thus supporting the establishment of new synaptic connections (**Figure 7**), including from inhibitory inputs from somatostatin interneurons, which are known to modulate PVs^28,92^. In this way, PV activity can be calibrated, allowing for IL firing during fear inhibition. By limiting PNN expression, PV INs gain an expanded dynamic range, allowing them to be active based on behavioral demands.

### Safety Conditioning Promotes Behavioral Flexibility

Our results indicate that safety conditioning has a distinct effect on remote memory by decreasing contextual fear over time, suggesting increased behavioral flexibility in the safety relative to the contextual fear conditioning group. Accordingly, despite both groups displaying similar levels of contextual freezing during recent retrieval, the SC group reduced defensive freezing at the remote timepoint whereas the CFC group did not. This reduction suggests that in the safety group, re-exposure to the context and the safety cue during recent retrieval, contributed to contextual fear extinction. However, the relative persistence of contextual fear in the CFC group, suggests a lack of flexibility, which is characteristic of the behavioral phenotype associated with PTSD^1,2,7,93–95^. Notably, safety conditioning also improves subsequent fear discrimination learning^7776,77^, indicating that safety conditioning can have long-term therapeutic effects on associative learning by helping increase behavioral flexibility^96^.

### Safety Conditioning Modulates PV activity and PNN Dynamics

The IL is known to be active during safety retrieval^12,13,97^, but the role of inhibition in sculpting IL activity during fear suppression is not yet well understood. Our data demonstrate that at recent retrieval, SC and CFC activate a similar proportion of PVs in the IL. At remote retrieval, in contrast, while the PV activity in the SC group remains relatively high, only very few PV INs are active in the CFC group. These data are consistent with previous reports on increased PV firing in the IL during threat avoidance, when PV inhibition in the IL impaired safety seeking^27,28^. Further, increased gamma power, which is associated with PV IN activity in the prefrontal cortex^30,98^, was observed in the IL during fear extinction^29^, a fear suppression behavior that is also highly dependent on the IL^18,24,99,100^. These effects may be due to increased IL activity overall, since PV cells are likely to be driven by local pyramidal cell firing^103^. On the other hand, chronic inhibition of GABA synthesis in the IL facilitated safety learning^11^ and chemogenetic inhibition of IL PV activity improved early extinction learning in males^26^, indicating that inhibition in the IL can also impair its function in safety learning. Approaches such as recordings and longitudinal tracking will help address this question in detail in the future, identifying the optimal level of PV activity for finetuning local cell firing and its effect on oscillations during behavior. Notably, our findings show that inhibiting PV INs during safety learning increases remote contextual fear and decreased neuro-glial interactions at remote retrieval, thus identifying longer-term impact of PV activity on behavioral and cellular changes.

Analogous to the pattern of PV activity described above, we detected similar levels of PNN surrounding PV INs in the fear and safety conditioning groups at recent retrieval, potentially reflecting a common early mechanism of learning-induced PNN assembly, linked to the recruitment of satOLIG2+ cells around the PV IN soma ^37,101,102^. However, at remote retrieval, only PV INs in the safety group were characterized by the presence of mature satOLIG2+ cells and significantly lower levels of PNNs compared to the CFC group. This suggests that while the initial learning phase may be similar in both groups, over time, only the CFC mice retain high levels of the PNN around PV IN, preserving the presynaptic inputs from learning^107^, thereby limiting plasticity and behavioral flexibility, and potentially stabilizing fear-related circuits. Notably, our results indicate that in the safety group, PV cells without PNNs showed a wider range of activity (20% active, 80% inactive), whereas those that had PNNs were 93% inactive. This suggests that safety conditioning may extend plasticity, allowing synaptic access to PV INs, which can excite or inhibit them, widening their dynamic range. It was previously shown that loss of PNN in the prelimbic region of the mPFC decreases PV input and increases glutamatergic input onto local pyramidal cells^107^, suggesting that similar changes may occur in the IL, which would help shift the overall balance to a more active IL. We propose that a balance in PV-PNN interactions is critical for flexible behavior: whereby PNNs stabilize memory circuits, but excessive PNNs may lead to inflexibility, limiting the capacity to update fear memories.

Further investigation of the molecular mechanisms underlying PNN remodeling and its effects on plasticity could uncover therapeutic targets for promoting behavioral flexibility.

## Methods

### Behavioral Protocol

#### Animals

PV-Cre Ai9 male mice (9-12 weeks, F1 offspring of homozygous C57B6J PV-Cre males (B6.129P2-*Pvalbtm1(cre)Arbr*/J, Jackson Labs) and homozygous Ai9-Cre females (B6.Cg-*Gt(ROSA)26Sortm9(CAG-tdTomato)Hze*/J, Jackson Labs), bred in house) were used for all experiments testing for cellular and myelin changes in SC and CFC groups. Male F1 PV-Cre mice (8–12 weeks, F1 offspring of homozygous male PV-Cre mice (B6.129P2-*Pvalbtm1(cre)Arbr*/J, Jackson Labs) and wild type female 129SvEv mice (Taconic)) were used for the optogenetic manipulation experiments.

Male 129SvEv mice (n=26, 9-12 weeks, Taconic BioSciences) were used for experiments testing RNA expression in safety vs fear conditioning (RNASeq,qPCR, RNAScope).

All mice were group-housed (3-4 per cage), provided ad libitum access to food and water, and kept in the Hunter College animal facility on a 12-hour light/dark cycle. Prior to the start of behavioral experiments, mice were single-housed (with bedding, dome shelters, and sunflower seeds) and handled for 2 days (5 min/day) before the start of the behavioral protocol. Animals were transported from the animal facility and acclimated in the lab for ∼ 1 hour before behavioral experiments were started each day. Mice were randomly assigned to SC, CFC or CA groups.

#### Behavioral Chamber

All behavior sessions were conducted in a chamber with grid floors (29.5 cm length x24.8 cm width x18.7 cm height, MedAssociates, St. Albans, VT). A house light (ENV-215 M 28 V, 100 mA, MedAssociates) was placed in the center of the chamber ceiling (18.7 cm above floor). The chamber was dimly lit (∼30 lux), and the house light briefly increased illumination in the chamber by 50-80 lux when turned on for 1 sec at the beginning of the CS presentations. Auditory cues (4 kHz) were delivered from a small speaker (ENV-224AM) embedded in a side panel of the chamber wall, 10.4 cm above the floor. Shocks (US) were delivered to the paws (0.6mA, 1 sec) with a constant current aversive stimulator (ENV-414S). A camera (Flex3, Optitrack, OR) was placed across from and slightly above the box to record each session for offline scoring of freezing. A blackout curtain separated the experimenter from the subjects and behavioral apparatus during the behavioral sessions. Chambers were cleaned with 100% ethanol between animals, and with non-scented soap and water at the end of each day.

#### Habituation

All groups underwent one day of habituation to the training context. During habituation, the SC and CA groups were presented with five cue trials (1-sec house light followed by a 30-sec, 4kHz, 50 ms pips, presented once a sec for 30 sec, with intertrial intervals [ITIs] ranging from 60-120 seconds. Cues began after ∼120 sec in the chamber, and sessions lasted 11 minutes on average. The CFC group was habituated to the chamber for 11 minutes.

#### Conditioning

Mice underwent two conditioning sessions, 24 hours after the last habituation session.

- **SC**: During each session, the SC group received five CS trials (1-sec house light co-occurring with a 30-sec tone), which was explicitly unpaired from five foot shock US (1-sec, 0.6mA) presentations. The mean inter-stimulus-interval (ISI) between any US and any proximal CS was 49sec (range 40-60sec), with some US trials without a CS trial in between, see Figure 1B). The inter-trial-interval (ITI) ranged from 60-120 sec.
- **CFC**: During each session, the CFC group received five unsignaled US trials (1-sec, 0.6mA), delivered at the same times within the session as in the SC group.
- **CA**: During each session the CA group received five CS trials (1-sec house light co-occurring with a 30-sec tone), delivered at the same times within the protocol as the cues in the SC group.

#### Recent and Remote Memory Retrieval

All animals underwent a recent retrieval session (24 hr after the second conditioning session), and mice in the remote retrieval session underwent a second retrieval session 21 days after the start of the experiment (18 days after second conditioning session). During each session:

- The **SC** and **CA groups** were presented with five CS trials of (1-sec house light co-occurring with a 30-sec tone) with a variable ITI of 60-120 sec.
- The **CFC group** was placed in the chamber for 15 minutes.

#### Conditioning and retrieval prior to transcriptional analysis

*RNAScope (Recent timepoint)*: To evaluate transcription of *Enpp6* in oligodendrocytes of the prefrontal cortex after fear or safety conditioning, mice were handled and habituated to the training context as described above. The SC (n=10) group was conditioned as described above. The fear group (n=9) underwent two sessions of classical fear conditioning, with five CS trials (1-sec house light co-occurring with a 30-sec tone) paired with the shock US (1-sec, 0.6mA) on each session (ITI 60-120 sec). Both groups underwent a recent retrieval session 24 h later, when they were placed back in the training context, and presented with 5 CS trials. Brain tissue was harvested after the recent retrieval session.

*RNASeq and qPCR (Remote timepoint):* To evaluate the transcription profile in the prefrontal cortex at remote timepoint after fear or safety conditioning, mice were handled, habituated to the training context, safety or fear conditioned, and underwent a recent retrieval session, as described above. Six days later, both groups underwent habituation to a testing context and two new tones (1kHz and 7kHz), and underwent three days of differential fear conditioning with the tones acting as a CS+ or CS- (1kHz and 7kHz tones, counterbalanced, ITI 60-120 sec). The CS+ was paired with a shock US (1-sec, 0.6mA) and the CS-was not paired with anything (as in Nahmoud et al., 2021^77^). The next day, both groups were placed in training context and exposed to five trials of each tone (10 trials total) for a differential fear conditioning retrieval session, and brain tissue was harvested the next day. The next day, mice were sacrificed for tissue collection for RNA sequencing (n = 3 FC, n = 4 SC) and qPCR analysis (n = 6 FC, n = 6 SC).

#### Behavioral Analysis

Custom-written scripts (Matlab, Mathworks) were used to cut all behavioral videos into 30-sec *context* and *cue* trials. For the CFC group, videos were cut at the identical times as the other groups, such that their context freezing was scored from the same times within the session as in the SC and CA groups (**Figure 1, Supplementary Figure 1**). The *Blind Analysis Tools* plugin (ImageJ^104^) was used to blind the experimenter to each subject’s group identify for manual scoring of the behavioral videos. A stopwatch was used to measure the defensive freezing (complete immobility except for breathing lasting ≥1 sec) during the context and the cue periods (expressed as a percentage of time in each trial). Context freezing is the average percent freezing during the 30 sec before trials 1 and 2 and cue-evoked freezing is the average percent freezing during trials 1-2.

#### Statistical Analysis

Prism (GraphPad) was used for statistical analysis and graphing, using grouped analyses, such as one-way analysis-of-variance (ANOVA) to evaluate percent freezing across behavioral groups, or a repeated measures (rm-)ANOVA to evaluate behavior in each group at different timepoints. Statistically significant main effects were further analyzed via multiple comparisons. Proportions were analyzed using Fisher’s Exact test. Data were visualized both as individual values via scatter plots, and as bars representing the group mean ±SEM.

### Perfusion and Tissue Collection

For cellular analysis, all animals were sacrificed 90-min after the start of either the recent or remote retrieval session to capture peak expression of the immediate early gene cFos, as a proxy for cellular activity. Mice were deeply anesthetized with an intraperitoneal (IP) injection of a mixture of ketamine (100mg/kg) and xylazine (10mg/kg), followed by thoracotomy and intracardiac perfusion with ice-cold 1X phosphate-buffered saline (PBS) and 4% paraformaldehyde (PFA) diluted in 1X PBS. Following fixation, animals were decapitated, brains were dissected and post-fixed in 4% PFA at 4°C for 24 hours in glass scintillation vials. Following post-fixation, brains were transferred to a solution of 30% sucrose in 1X PBS with 2% sodium azide (NaAz) for cryoprotection for ∼48-72 hours (or until brains sunk to the bottom of the vial). Brains were then embedded in OCT-filled plastic molds and flash-frozen using 2-methylbutane and dry ice. Brains acclimated in a cryostat (Cryostar, Leica) for ∼20 minutes before sectioning at 14 μm. Tissue sections were stored in light-sensitive well plates in 1X PBS with NaAz at 4°C until staining.

### Immunohistochemistry

Coronal brain slices containing the medial prefrontal cortex (mPFC; ∼Bregma +1.40 to +1.9 mm) were mounted onto SuperFrost Plus microscope slides. Immunohistochemistry (IHC) was performed on slide-mounted tissue, encased in a hydrophobic barrier created with Kwik-Cast Silicon Elastomer and stored in humidity chambers throughout staining. The general IHC protocol consisted of three washes in 1X PBS (five minutes each), followed by blocking for one hour at room temperature in blocking solution (10% normal donkey serum (NDS, Jackson Immuno Research, 017-000-121) in PBST (1X PBS with 0.3% Triton-X)). Tissue was then incubated overnight at 4°C in the primary antibody solution, diluted in blocking solution. After 24 hours, the tissue was washed again three times with PBS and incubated in secondary antibody for one hour at room temperature. Following secondary antibody incubation, the tissue was washed once with PBST for ten minutes and twice with PBS for five minutes each. The hydrophobic barriers were then removed, and the slides were cleaned and coverslipped with Prolong Diamond Mounting Reagent with DAPI (ThermoFisher Scientific, P36971).

For cFos detection, tissue was incubated with a rabbit anti-cFos primary antibody (1:2000, Abcam,190289) and donkey anti-rabbit Alexa Fluor 488 (1:500, Life Technologies, AF21206) as the secondary antibody, both diluted in blocking solution. We identified PNNs using a chemical stain for wisteria fluorobunda agglutinin (WFA, Sigma, L1516), which is a lectin that has a high affinity for terminal N-acetylgalactosamine residues within the PNN^105^. Biotinylated WFA was added to the primary antibody solution (1:200) and visualized using fluorescently tagged streptavidin for signal amplification, which was included in the secondary antibody solution. PV INs were identified via endogenous red fluorescence (tdTomato, pseudo-colored cyan in the figures). See Table 1 for complete information regarding antibodies used throughout this manuscript.

### Microscopy

An SP8 (Leica) confocal microscope was used to image immuno-stained tissue. Fluorophores for Alexa Fluor 488, 647, DyLight 405 (see Table 1 for information), and tdTomato were visualized using line sequential scanning to prevent crosstalk. Imaging parameters included a 524 x 524-pixel resolution, a scan speed of 600, a 40x objective lens with 0.75x zoom, and line averaging (2 lines). Images were acquired as 3D Z-projections consisting of 10 optical slices, with 24-tile scans for coverage. Two slices of the mPFC were taken per animal using identical settings throughout all cohorts.

### Image Pre-Processing and Analysis

Raw image files (lif format) were imported into ImageJ ^104^ . A custom protocol was used to align images to the Allen Brain Atlas at the corresponding Bregma depth (∼1.70 mm) using known measurements corresponding to landmarks like the dorsal surface, midline, and corpus callosum. Regions of interest (ROIs) corresponding to the prelimbic (PL) and infralimbic (IL) cortices were selected, cut from the rest of the tissue, measured for area in mm^2^, and saved as new files (tiff format). The *Blind Analysis Tools* plugin (ImageJ^104^) was used to randomize file names so as to blind the experimenter to group identity, and were then imported into Imaris software (10.1.1, Oxford Instruments) for analysis.

During image pre-processing (Imaris), Gaussian filtering and background subtraction were consistently applied across all stains. The *Surface* tool was used for object detection of PV, PNNs, and cFos markers and a batch process was used to uniformly apply surface generation settings. Surfaces were manually adjusted post-processing, and raw cell counts were recorded. Additional surfaces were created to for overlap analyses (e.g., PV cells with cFos, PV cells with PNNs). Co-localizing structures were defined as those with zero distance (in µm) between different surface categories for each stain.

#### Myelination

Global myelination was quantified by calculating the Corrected Total Myelin Fluorescence (CTMF) from IHC-labeled myelin basic protein (MBP). CTMF was adapted from a Corrected Total Cell Fluorescence (CTCF) protocol ^106^. CTMF was determined using the following equation:

*CTMF = Integrated Fluorescence Density of MBP – (Area of ROI * Mean Fluorescence of Background)*

#### OLC Identification

IHC stains for OL lineage markers were used as a proxy of OLC maturation status. An Olig2 primary antibody was used as a marker for all oligodendroglial cells across the lineage (i.e., OLCs). OLCs were defined as cells co-expressing Olig2 and DAPI (i.e., Olig2^+^:DAPI^+^) and the density of OLCs was calculated using the following equation:

*cells/mm^2^ = Total Olig2^+^:DAPI^+^ cells ÷ Area (mm^2^)*

OLs were identified with Olig2, DAPI, and the CC1 antibody, which recognizes the adenomatous polyposis coli (APC) protein which is highly expressed in differentiated cells (Olig2^+^:CC1^+^:DAPI^+^). OPCs were initially identified using the neural/glial antigen 2 (NG2) antibody. However, preliminary staining trials showed that all NG2+ OPCs were also CC1-, and we discontinued using the NG2 stain, to maximize fluorophore availability. Therefore, OPCs were identified as Olig2^+^:CC1^-^:DAPI^+^ cells.

The maturation status of OLCs was measured using the following equation:

*% of OLC population that is an OPC or OL = # of OPCs (or OLs) ÷ Total # of OLCs * 100*

#### sat*Olig2^+^* cells

Sat*Olig2^+^* cells were identified as any Olig2^+^:DAPI^+^ OLC in direct contact with the soma of a PV IN. Sat*Olig2^+^*-PV pairs were quantified as a percentage of the PV IN population by using the following equation:

*% PV-* sat*Olig2^+^ pairs = # of PV^+^:Olig2^+^:DAPI^+^ cells ÷ Total # of PV^+^:DAPI^+^ cells * 100*

Sat*Olig2^+^*-PV pairs surrounded by PNNs were quantified as a percentage of the total population of PV- sat*Olig2^+^* pairs by using the following equation:

*% PV-* sat*Olig2^+^cells with PNN = PV^+^:Olig2^+^:WFA^+^ cells ÷ PV^+^:Olig2^+^cells * 100*

#### Data Analysis and Statistics

Cell counts were recorded in Excel, and the image file names were unblinded after all analyses were complete. Prism (GraphPad) was used for statistical analysis.

### Bulk RNAseq, qPCR, RNAScope

For qPCR and RNASeq, tissue from safety and fear-conditioned mice was collec13 days post-conditioning (see *Behavior* section). After anesthesia (dry ice) and cervical dislocation, brains were dissected, and the mPFC was isolated and stored at -80°C. Total RNA was extracted and processed for bulk RNA sequencing (RNA-seq) at the Epigenetics Facility (ASRC, CUNY). Quality of the RNA was assessed by Tapestation 4200 (Agilent Technologies). Only RNA with RIN>=8 was used for subsequent library construction. mRNA-seq libraries were prepared using KAPA mRNA Hyperprep kit (Roche). Briefly, mRNA was captured using oligo-dT beads, followed by random fragmentation. First strand cDNA was synthesized using mRNA template and random hexamer primers, after which second-strand cDNA was synthesized combined with A-tailing. After a series of terminal repair, ligation and sequencing adaptor ligation, the double stranded cDNA library was completed through size selection and PCR enrichment. The libraries were sequenced on Illumina NovaSeq. Depths of minimum 30 million paired-end 150 bp reads were generated for each sample. Entire dataset was deposited into GEO data depositary (accession number GSE285736). Pathway analysis identified enriched biological pathways based on differentially expressed gene profiles. For qPCR, cDNA was synthesized, and gene expression of select OL and myelin-related genes (e.g., *Myrf*, *Enpp6*, *Mbp*, *Mobp*) was quantified.

For RNAScope, cryosectioned tissue from perfused mice, was mounted onto slides and probes were used for *Enpp6*, combined with IHC for Olig2 and counterstaining with DAPI. To identify *Enpp6* gene expression in OLs, colocalization of *Enpp6* transcripts by OLCs was analyzed.

### Single-cell RNA sequencing Data Mining

scRNAseq (10x) data was obtained from the Allen Institute’s open-source knowledge base, *ABC Atlas* (https://knowledge.brain-map.org/abcatlas). Filters were set for sex, (males), dissection region (PL-IL-Orb, i.e., mPFC), genes of interest, and cell type (subdivided by class, subclass, and supertype). In particular, the OPC-Oligo class (#31) is subdivided into OPC (#326) and OL (#327) subclasses, with a focus on OL supertypes: committed-OLs (COPs, # 1181), NFOLs (#1182), MFOLs (#1183), and MOLs (#1184). Gene expression percentages were calculated by importing ABC Atlas UMAP images into ImageJ, measuring expression density (cells/mm²) within each supertype, and normalizing to the total supertype density. UMAP images were converted to 8-bit, thresholded, and a selection was created to measure gene expression density (cells/mm²), which was reported as a percentage of total baseline gene expression within the OLC supertype (without gene filtering).

### Stereotaxic Surgery

For the optogenetic inhibition experiments, PV-Cre mice underwent a stereotactic surgery for virus infusion and optic fiber implantation. For all surgeries, mice were anesthetized with 2% isoflurane in oxygen, placed in a stereotaxic frame (Kopf Instruments, Tujunga, CA) and maintained on 1.5% isoflurane throughout surgery (oxygen at a flow rate of 1L/min). Temperature was maintained at 36C±1C with a feedback-regulated heating pad. Mice received dexamethasone (1mg/mL, s.c.) and bupivacaine under the scalp (5mg/mL, s.c.) prior to incision. Craniotomies were performed bilaterally at the injection site using a drill with a burr attachment, and the viral vector (0.3 µL per hemisphere) was injected at a rate of 0.08μL/min using a 10μL Hamilton syringe (QSW Stereotax Injector, Stoelting, IL). Following the injection, the syringe was left in place for an additional 6 minutes, pulled up 1mm and left another 5 minutes to allow for viral diffusion before being slowly withdrawn.

#### Viral Injection

Using PV-Cre mice, we selectively infected PV INs of the IL (AP, +1.8mm, ML, ±0.4 mm DV, -2.4mm from brain surface) with a Cre-dependent adeno-associated viral vector (AAV5) encoding for enhanced archaeorhodopsin 3.0 (eArch) fused with enhanced yellow fluorescent protein (rAA5-Ef1a-DIO-eArch3.0-EYFP). The control group received a virus that only expressed eYFP (rAA5-Ef1a-DIO-eArch3.0-EYFP

#### Optical Fiber Implantation

Optic fibers (200 µm core diameter, 0.37 NA, Thorlabs) were implanted bilaterally 0.1 mm above the virus injection sites in the IL (AP +1.8 mm, ML ±0.4 mm, DV -2.3 mm from brain surface). Fibers were secured with Relyx U200 self-adhesive UV-curing resin cement (3M ESPE) curing solution, as well as Metabond (Parkell) and Orthojet (Lang Dental) dental cement.

#### Postoperative Care

At the completion of the surgery, mice were given carprofen (5mg/kg) for analgesia and monitored for recovery from surgery. They were given 3 weeks to recover from surgery and to ensure stable viral expression before any beginning behavioral experiments.

### Optogenetics Behavioral Protocol

Both groups of mice (eArch and eYFP) underwent safety conditioning. First, mice were habituated to the training context and the safety cue (5 trials, as described above), and then in a separate habituation session, mice received 5 trials (523 nm, 10±1mW, 40 sec, 5-sec ramp modulated on and 5-sec off) of laser exposure. Both groups underwent SC protocols as described above. During safety conditioning, the laser was delivered bilaterally during each safety CS, starting 5 seconds before and ending 5 seconds after the safety cue (523 nm, 10±1mW, 40 sec, 5-sec ramp modulated on and 5-sec off). Both groups underwent Recent Retrieval (24 hours) and Remote Retrieval (21 days) where they were re-exposed to the training context and the safety CS (five, 30-sec trials). The laser was not turned on during retrieval sessions. All behavior was recorded by a video camera and analyzed offline for defensive freezing by an observed blinded to group. For optogenetics behavior data plotted in **Figure 6** and **Supplementary Figure 5**, “context” measurements are the average freezing (%) during “context” pre-trials 1 and 2 and “cue” measurements are the average freezing (%) during cue trials 1-5, calculated by the equation: # seconds spent freezing/30 sec [trial length]/100.

## Author contribution

L.E.D. conducted all experiments, analyzed data, contributed to the study design, and wrote the manuscript. J.L. performed experiments, analyzed data, and edited the manuscript. I.N. conducted experiments. P.C. designed and supervised the study, and edited the manuscript, E.L. designed and supervised the study, and wrote the manuscript.

## Acknowledgments

We thank Dr. Carmen Melendez-Vasquez for valuable discussion and guidance on data interpretation and analysis throughout this project, and Tikva Nabatian for assistance with behavioral experiments. This work was funded by the ASRC Seed grant (E.L. and J.L.), the GC Provost’s Seed Grant (E.L. and P.C.), NIH-NIMH R01MH118441 (E.L.), NIH-NINDS R35 NS111604 (P.C.).

## Supplementary

**Supplementary Figure 1. Cue alone (CA) behavior during recent and remote retrieval. (A)** Sample conditioning session. CFC group (top, orange) received five foot shocks. SC (middle, blue) received five foot shocks US unpaired in time with five cue CS. CA (bottom, purple) received five cue presentations without any shocks US. **(B)** At recent retrieval, the CA group shows no differences in percent freezing between context and cue (paired t-test, p=0.65). **(C)** At remote retrieval, the CA group also shows no differences in percent freezing between context and cue (paired t-test, p=0.74). **(D)** At recent retrieval, the CA group is freezing less to context than the CFC and the SC groups (one-way ANOVA F(2,22)=10.17, p=0.0007, CFC vs SC p=0.23, CFC vs CA, p=0.005, SC vs CA p=0.006). **(E)** At remote retrieval, only the CFC group shows higher contextual fear than the CA group (one-way ANOVA F(2,10)=7.166, p=0.012, CFC vs CA p=0.009, SC vs CA p=0.15). **(F)** Percent contextual freezing at recent and remote retrieval in the CA group is similarly low (paired t-test, p=0.4). Bars represent mean ± SEM.

**Supplementary Figure 2. Activity profiles of PV INs with and without PNNs. (A)** PV INs with a PNN are generally inactive (PV:WFA+:cFos+/ PV:WFA) during recent and remote retrieval in SC and CFC groups. **(B)** At remote safety retrieval, a higher proportion of PV cells without PNNs are active (cFos+) compared to PVs with PNNs (Χ^2^, Fisher’s exact test, p=0.049). **(C)** PV cells without PNNs are overall more inactive (cFos-) than active (Unpaired t-test p=0.02).

**Supplementary Figure 3. At recent and remote retrieval, there is higher mPFC transcription of myelin and oligodendrocyte-related genes after safety than fear conditioning. (A)** Experimental timeline for testing oligodendrocyte gene transcription at recent retrieval. On Day 1-2, mice were habituated to the training context and the 4kHz cue (50ms pips, presented at 1Hz for 30 sec), and then on Days 3-4 underwent 2 days of either fear conditioning (FC, CS-US paired) or safety conditioning (SC, CS-US unpaired), followed by a recent retrieval on Day 5 when mice were placed back in the training context and were exposed to five presentation of the cue CS. (**B)** Examples of RNAScope staining, showing *Olig2* (magenta) and *Enpp6* expression (green dots) at recent fear retrieval (top) and safety retrieval (bottom). Nuclei are counterstained with DAPI (blue). Arrowheads show the overlap of *Enpp6+/*OLIG2+ in OLs. **(C)** Significant increase of *Enpp6+/*OLIG2+ NFOLs in safety (blue, n=3 mice) compared to fear (red, n=3 mice) trained animals (Mann-Whitney, p<0.01). **(D)** Experimental timeline for testing oligodendrocyte and myelin gene transcription at remote retrieval. On Day 1-2, mice were habituated to the training context and the 4kHz cue (50ms pips, presented at 1Hz for 30 sec), and then on Days 3-4 underwent 2 days of either fear conditioning (FC, CS-US paired) or safety conditioning (SC, CS-US unpaired), followed by a recent retrieval on Day 5 when mice were placed back in the training context and were exposed to five presentation of the cue CS. Then, on Days 10-12, mice underwent differential fear conditioning with two new tones (2kHz, 8kHz tones) and then, the next day (Day 13), underwent differential fear retrieval in a new context. On Day 14, the mPFC was collected for bulk RNASeq or for targeted qPCR analysis. (**E)** qPCR in mice (n=6/grp) show increased transcription of myelin related genes in safety relative to fear trained mice, two-way ANOVA (F(1,60)=14.79, p<0.001). Data presented as mean ± SEM. **(F)** Heatmap of the top 20-most differentially expressed transcripts in the mPFC shown by subject. Overall, fear (n=3, red) vs safety (n=4, blue) trained subjects cluster together forming separate clusters. Green indicates higher and yellow indicates lower gene expression. **(G)** Enriched pathway analysis showing the top ontology terms of differentially expressed genes between safety and fear trained animals. Note that the top two pathways that differentiate the two groups contain axon ensheathment and myelination. **(H)** Principal Component Analysis (PCA) showing the overall segregation in transcriptome profiles along the two first principal components of animals who were trained on safety (blue) or fear (red) conditioning. **(I)** Volcano plot showing the fold change in upregulated genes of safety trained animals, red indicates significantly upregulated and downregulated genes.

**Supplementary Figure 4. Overall myelin content in the IL doesn’t change between CFC and SC. (A)** Experimental timeline for immunohistochemical analysis to measure myelin content at recent and remote retrieval of safety or contextual fear conditioning. **(B)** *Left,* Example of myelin basic protein (MBP, green) staining in the mPFC. *Right top,* Example of background (1) and signal (2) ROI selections used for intensity measurement. *Right bottom*, expansion of background and signal. **(C)** Corrected total myelin fluorescence (CTMF) is the same in the CFC and SC groups at recent retrieval and remote retrieval (two-way ANOVA: group F(1,3) = 1.82, p=0.27, timepoint F(1,4) = 4.12, p=0.11, group x timepoint F(1,3)=1.52, p=0.31). CTMF: (integrated density in ROI) – [(area of ROI) * (Background mean intensity)].

**Supplementary Figure 5. Optogenetic inhibition of IL PV INs during conditioned safety. (A)** Percent freezing to context and cue at remote retrieval after PV inhibition during safety learning. The eYFP group shows similar levels of freezing whereas the eArch group shows decreased freezing during the cue relative to context (two-way ANOVA: trial type, F(1,12)=5.34, p=0.039; multiple comparisons: eYFP context vs cue p=0.27, eArch context vs cue p=0.049. **(B)** Density of IL PV cells expressing virus is the same in the eYFP and eArch groups (unpaired t-test p=0.15).

**Table 1.** Antibodies used for immunohistochemistry.

