## Supplemental Figures and Table for "PV - Oligodendrocyte Interactions in the Infralimbic Cortex Promote Extracellular Plasticity after Safety Learning"

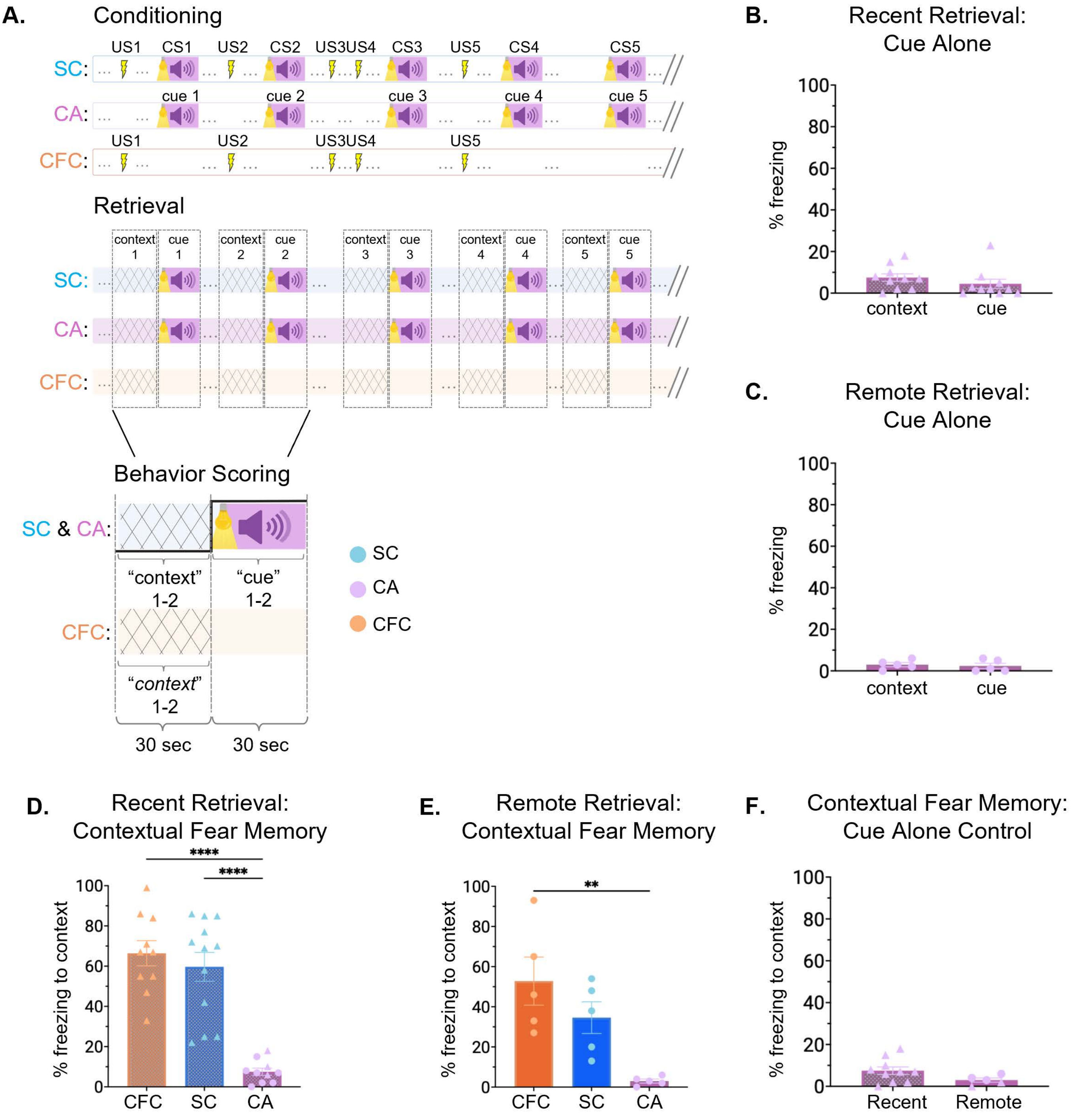


**Supplementary Figure 1. Cue alone (CA) behavior during recent and remote retrieval. (A)** Sample conditioning session. CFC group (top, orange) received five foot shocks. SC (middle, blue) received five foot shocks US unpaired in time with five cue CS. CA (bottom, purple) received five cue presentations without any shocks US. **(B)** At recent retrieval, the CA group shows no differences in percent freezing between context and cue (paired t-test, $p=0.65$). **(C)** At remote retrieval, the CA group also shows no differences in percent freezing between context and cue (paired t-test, $p=0.74$). **(D)** At recent retrieval, the CA group is freezing less to context than the CFC and the SC groups (one-way ANOVA $F(2,22)=10.17$, $p=0.0007$, CFC vs SC $p=0.23$, CFC vs CA, $p=0.005$, SC vs CA $p=0.006$). **(E)** At remote retrieval, only the CFC group shows higher contextual fear than the CA group (one-way ANOVA $F(2,10)=7.166$, $p=0.012$, CFC vs CA $p=0.009$, SC vs CA $p=0.15$). **(F)** Percent contextual freezing at recent and remote retrieval in the CA group is similarly low (paired t-test, $p=0.4$). Bars represent mean ± SEM.

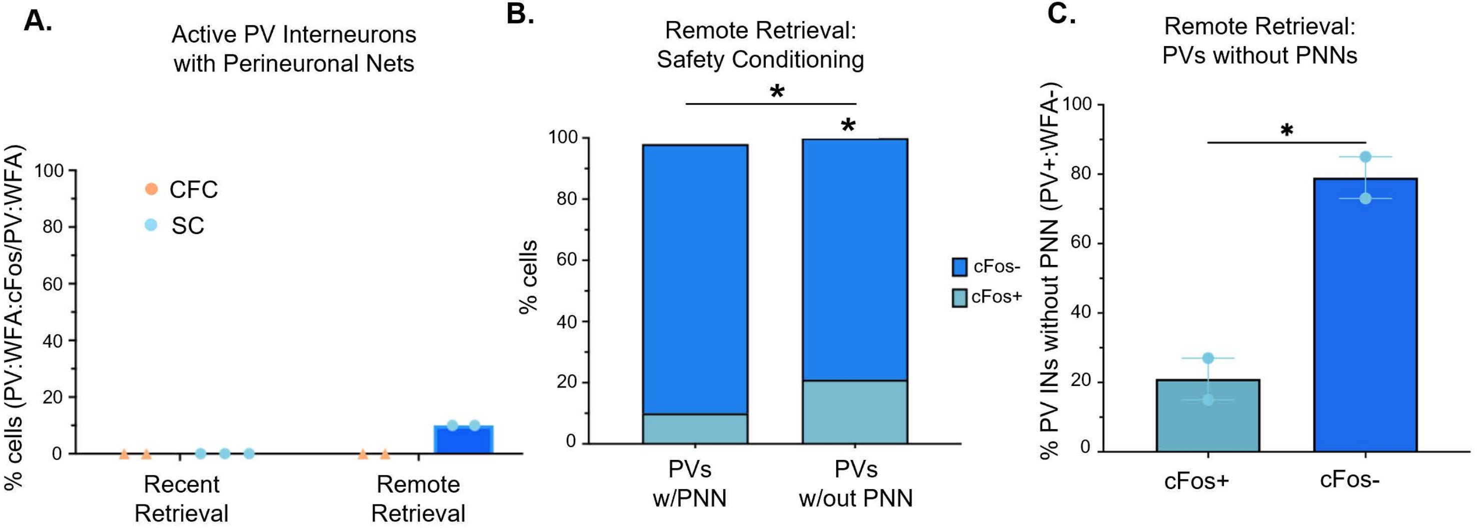


**Supplementary Figure 2. Activity profiles of PV INs with and without PNNs. (A)** PV INs with a PNN are generally inactive (PV:WFA+:cFos+/ PV:WFA) during recent and remote retrieval in SC and CFC groups. **(B)** At remote safety retrieval, a higher proportion of PV cells without PNNs are active (cFos+) compared to PVs with PNNs ($X^2$, Fisher's exact test, p=0.049). **(C)** PV cells without PNNs are overall more inactive (cFos-) than active (Unpaired t-test p=0.02).

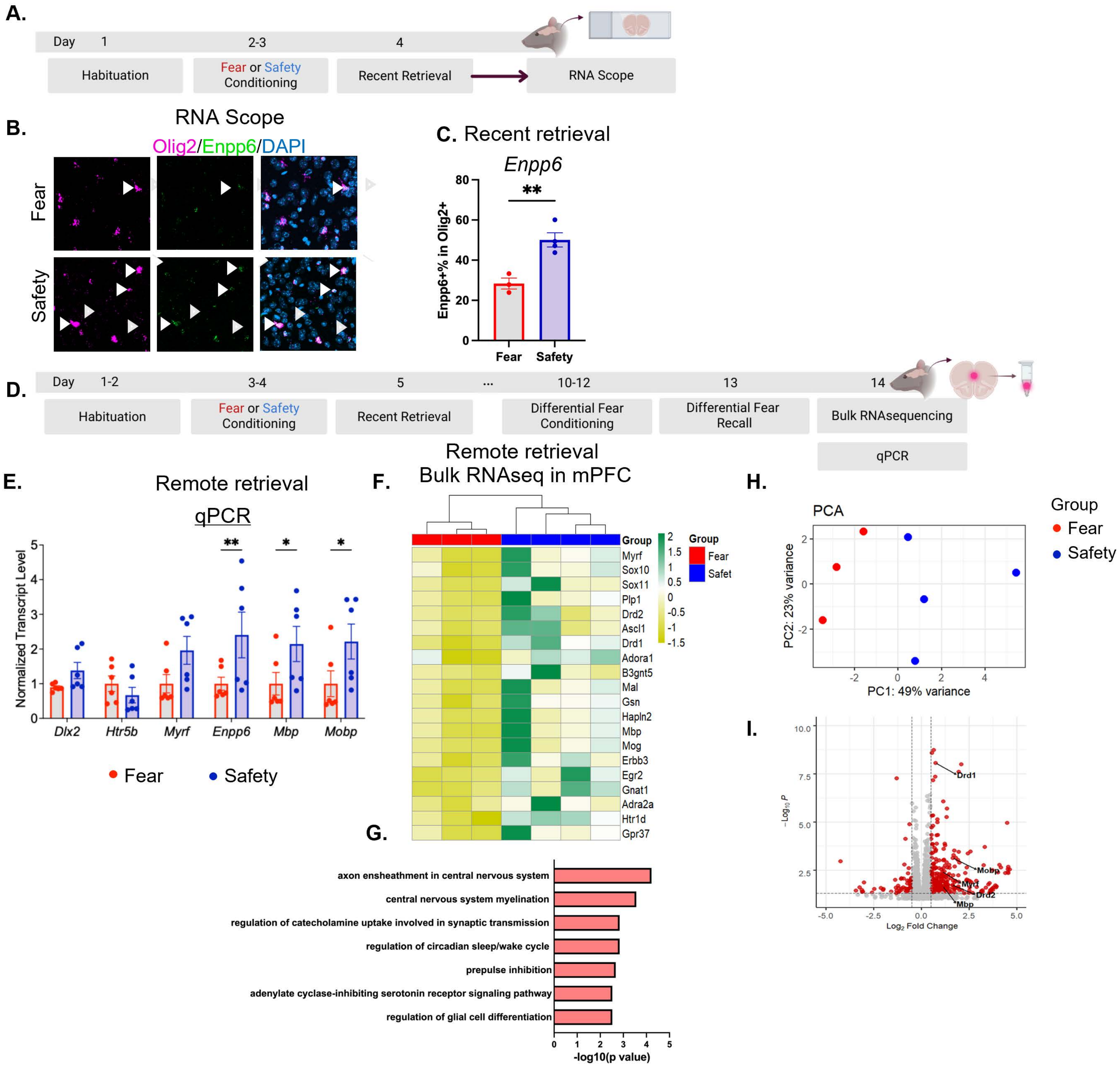


**Supplementary Figure 3. At recent and remote retrieval, there is higher mPFC transcription of myelin and oligodendrocyte-related genes after safety than fear conditioning. (A)** Experimental timeline for testing oligodendrocyte gene transcription at recent retrieval. On Day 1-2, mice were habituated to the training context and the 4kHz cue (50ms pips, presented at 1Hz for 30 sec), and then on Days 3-4 underwent 2 days of either fear conditioning (FC, CS-US paired) or safety conditioning (SC, CS-US unpaired), followed by a recent retrieval on Day 5 when mice were placed back in the training context and were exposed to five presentation of the cue CS. (**B**) Examples of RNAScope staining, showing *Olig2* (magenta) and *Enpp6* expression (green dots) at recent fear retrieval (top) and safety retrieval (bottom). Nuclei are counterstained with DAPI (blue). Arrowheads show the overlap of *Enpp6+/*OLIG2+ in OLs. **(C)** Significant increase of *Enpp6+/*OLIG2+ NFOLs in safety (blue, n=3 mice) compared to fear (red, n=3 mice) trained animals (Mann-Whitney, $p<0.01$). **(D)** Experimental timeline for testing oligodendrocyte and myelin gene transcription at remote retrieval. On Day 1-2, mice were habituated to the training context and the 4kHz cue (50ms pips, presented at 1Hz for 30 sec), and then on Days 3-4 underwent 2 days of either fear conditioning (FC, CS-US paired) or safety conditioning (SC, CS-US unpaired), followed by a recent retrieval on Day 5 when mice were placed back in the training context and were exposed to five presentation of the cue CS. Then, on Days 10-12, mice underwent differential fear conditioning with two new tones (2kHz, 8kHz tones) and then, the next day (Day 13), underwent differential fear retrieval in a new context. On Day 14, the mPFC was collected for bulk RNASeq or for targeted qPCR analysis. (**E)** qPCR in mice (n=6/grp) show increased transcription of myelin related genes in safety relative to fear trained mice, two-way ANOVA ($F(1,60)=14.79$, $p<0.001$). Data presented as mean ± SEM. **(F)** Heatmap of the top 20-most differentially expressed transcripts in the mPFC shown by subject. Overall, fear (n=3, red) vs safety (n=4, blue) trained subjects cluster together forming separate clusters. Green indicates higher and yellow indicates lower gene expression. **(G)** Enriched pathway analysis showing the top ontology terms of differentially expressed genes between safety and fear trained animals. Note that the top two pathways that differentiate the two groups contain axon ensheathment and myelination. **(H)** Principal Component Analysis (PCA) showing the overall segregation in transcriptome profiles along the two first principal components of animals who were trained on safety (blue) or fear (red) conditioning. **(I)** Volcano plot showing the fold change in upregulated genes of safety trained animals, red indicates significantly upregulated and downregulated genes.

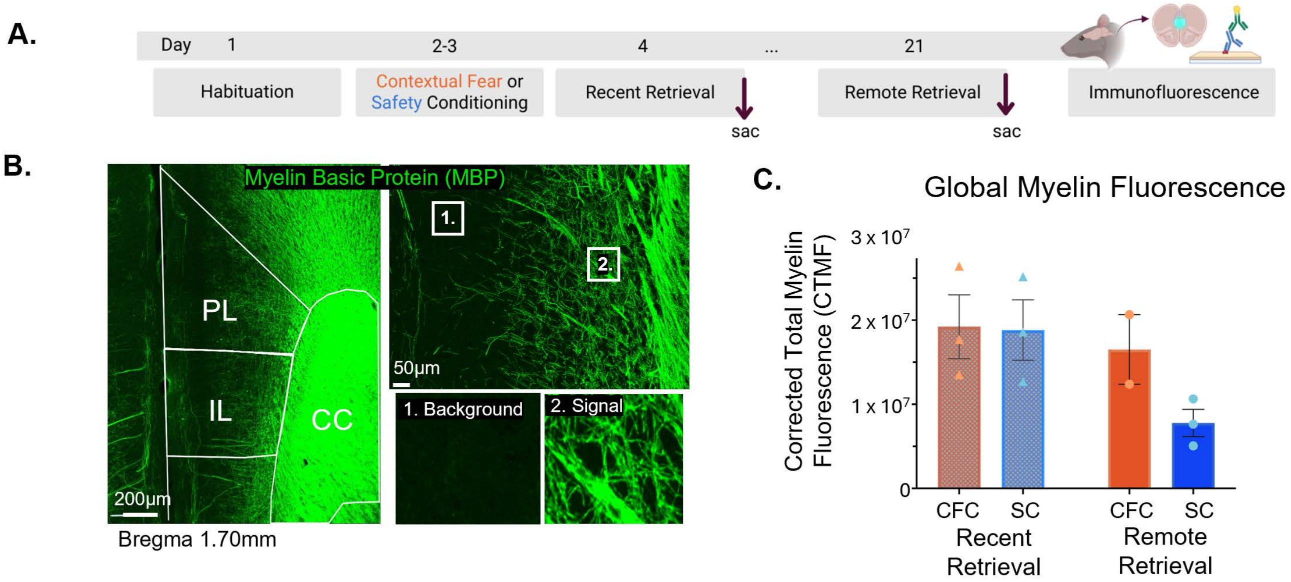


**Supplementary Figure 4. Overall myelin content in the IL doesn't change between CFC and SC. (A)** Experimental timeline for immunohistochemical analysis to measure myelin content at recent and remote retrieval of safety or contextual fear conditioning. **(B)** *Left,* Example of myelin basic protein (MBP, green) staining in the mPFC. *Right top,* Example of background (1) and signal (2) ROI selections used for intensity measurement. *Right bottom*, expansion of background and signal. **(C)** Corrected total myelin fluorescence (CTMF) is the same in the CFC and SC groups at recent retrieval and remote retrieval (two-way ANOVA: group $F(1,3) = 1.82$, $p=0.27$, timepoint $F(1,4) = 4.12$, $p=0.11$, group x timepoint $F(1,3)=1.52$, $p=0.31$). CTMF: (integrated density in ROI) – [(area of ROI) * (Background mean intensity)].

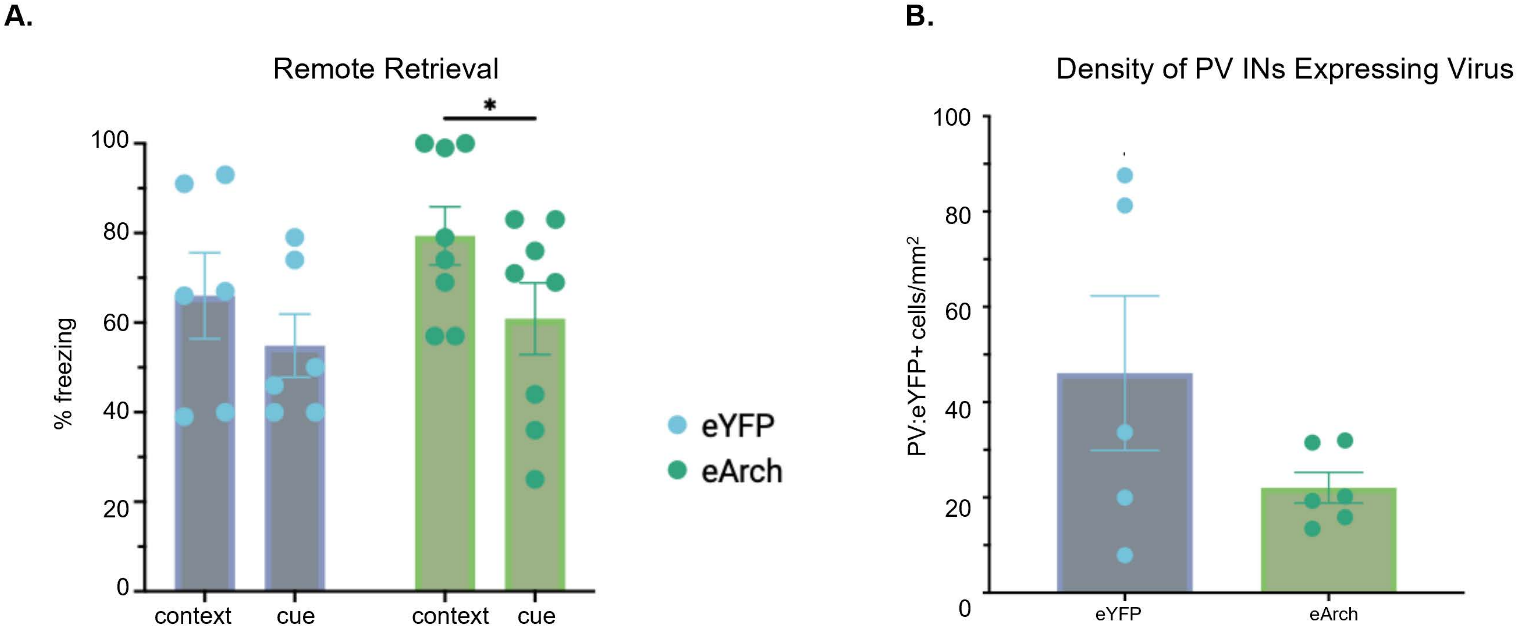


**Supplementary Figure 5. Optogenetic inhibition of IL PV INs during conditioned safety. (A)** Percent freezing to context and cue at remote retrieval after PV inhibition during safety learning. The eYFP group shows similar levels of freezing whereas the eArch group shows decreased freezing during the cue relative to context (two-way ANOVA: trial type, $F(1,12)=5.34$, $p=0.039$; multiple comparisons: eYFP context vs cue $p=0.27$, eArch context vs cue $p=0.049$. **(B)** Density of IL PV cells expressing virus is the same in the eYFP and eArch groups (unpaired t-test $p=0.15$).

| Anti | Host | Dilution | Company / ID | Anti | Host | Label | Dilution | Company / ID |
|---|---|---|---|---|---|---|---|---|
| PV | Guinea Pig | 1:500 | Synaptic Systems 195-004 | Guinea Pig | Donkey | DyLight 405 | 1:200 | Jackson 706-475-148 |
| CC1 | Mouse | 1:500 | Abcam ab16794 | Mouse | Donkey | AF 594 | 1:500 | Jackson 715-585-150 |
| Olig2 | Rabbit | 1:1000 | Millipore AB9610 | Mouse | Doney | AF 488 | 1:500 | Southern Biotech 6415-30 |
| Olig2 | Mouse | 1:200 | Millipore MABN50 | Rabbit | Donkey | AF 647 | 1:500 | Thermo Fisher A31573 |
| cFos | Rabbit | 1:2000 | Abcam 190289 | Rabbit | Donkey | AF 488 | 1:500 | Life Technologies A21206 |
| NG2 | Rabbit | 1:200 | Millipore AB5320 | Streptavidin | *N/A* | AF 647 | 1:1000 | Jackson 016-600-084 |
| MBP | Mouse | 1:500 | BioLegend 808401 | Streptavidin | *N/A* | DyLight 405 | 1:500 | Thermo Fisher 21831 |
| WFA | *N/A* | 1:200 | Sigma L1516 | | | | | |

**Table 1:** Antibodies used for immunohistochemistry.
